# Exosomal microRNA signature from plasma-derived extracellular vesicles in gastric cancer

**DOI:** 10.1101/2023.04.28.538562

**Authors:** Andrés Rincón-Riveros, Victoria E. Villegas, Nicolle Stefania Quintero Motta, Liliana López-Kleine, Josefa Antonia Rodríguezand

## Abstract

**Background:** Gastric cancer is a heterogeneous pathology that represents the fifth most frequent malignancy in the world, with more than 750,000 deaths by 2020. With significant repercussions in public health, this pathology lacks biomarkers for early diagnosis, with endoscopy biopsy being the golden test for its detection. In the exploration of new strategies to control gastric cancer in recent years, liquid biopsy appears as a potential source of biomarkers using non-invasive procedures.

**Methods:** Here we present the characterization of miRNAs contained in plasma-derived exosomes from patients with gastric cancer. Extracellular vesicles (EVs) were isolated using size-exclusion chromatography (SEC) and their characterization was performed by electron microscopy, protein expression, and nanoparticle analysis techniques. Total RNA from isolated exosomes was obtained for small RNA-seq analysis.

**Results:** Transcriptomic miRNA data were used to identify differentially expressed miRNAs between patients with benign and malignant gastric diseases, which resulted in a molecular signature of nine miRNAs, that were used in a regression model to classify individuals as either having benign or malignant disease. Further, we compared benign-malignant patients at different stages of gastric cancer, and we detected 15 differentially expressed miRNAs. Among these 15 miRNAs, miR-92a-3p, miR451a, and miR126-3p were identified as winners due to their clinical and regulatory relevance.

**Results:** Our results offer relevant information of a Colombian case study allowing us to propose three transcriptomic gastric cancer biomarkers in liquid biopsy.

**Graphical Abstract:** 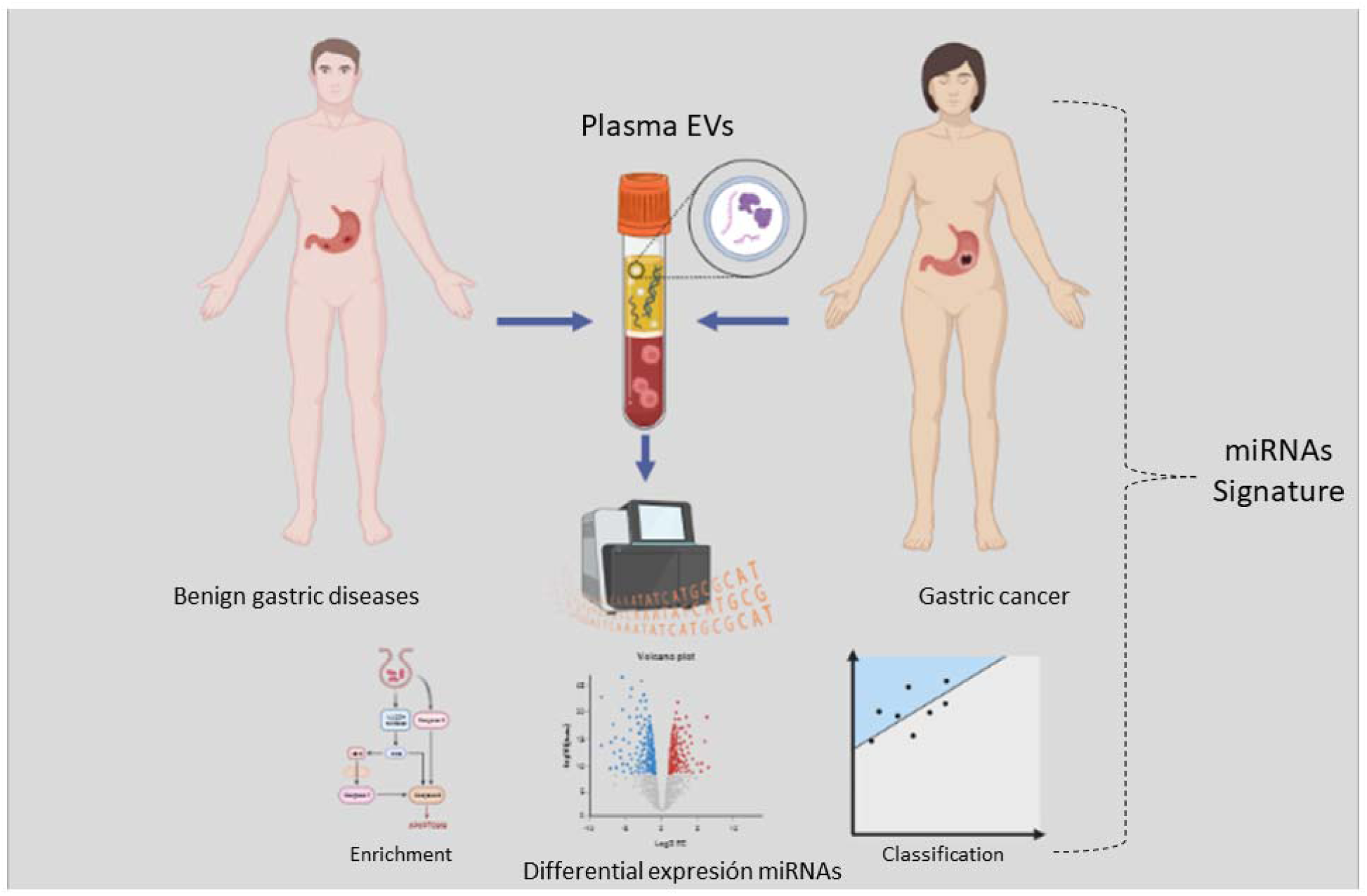

**Summary:** EVs are structures surrounded by a lipid bilayer that facilitate intercellular communication by transporting biomolecules commonly referred to as part of liquid biopsy. In this study, we examine the miRNAs contents of plasma isolated EVs from patients with both benign gastric diseases and gastric cancer to identify potential biomarkers for gastric cancer.

## Introduction

Gastric cancer is one of the more incident type of cancer with high incidence and mortality worldwide, which ranks fifth among incidents with more than 1 million new cases which corresponds to 6% of total cancer worldwide by 2020 (GLOBOCAN). Mortality exceeds 750,000 deaths per year. Gastric cancer is 50% more common in men than in women. According to Cancer Today of the International Agency for Research on Cancer (IARC), in Colombia, gastric cancer is the leading cause of cancer death in men, with approximately 3,963 deaths, and the fourth cause of death in women with more than 2,400 deaths per year [1]. Gastric cancer is considered a public health problem given its high mortality, due in part to the difficulty of making a timely diagnosis. Consequently, 5-year survival is less than 30%. The golden test for the detection of this type of cancer is endoscopy with biopsy, but it is not cost-effective as a screening test in low- or middle-income countries [2].

Currently explored new diagnostic strategies may lead to the development of non-invasive methodologies for early diagnosis and monitoring of this cancer [3] such as liquid biopsy, which emerges as a tool to search for biomarkers with clinical utility in diagnosis and disease monitoring or to obtain information that allows an adequate choice of personalized therapies in real time. Liquid biopsy allows the evaluation of circulating tumor cells (CTCs), circulating nucleic acids, proteins, and EVs, among others [4].

EVs are cellular structures surrounded by a lipid bilayer, secreted by most nucleated cells and bacteria. They can be classified into apoptotic bodies, microvesicles, and exosomes, based on their size. Exosomes are EVs with an average diameter of 30 to 150 nm [5]. They are found in biological fluids and perform normal physiological processes such as intercellular communication. However, in pathological conditions such as cancer, they can promote tumor invasion and metastasis. Recently, there has been much interest in the study of exosomes because they are carriers of molecules such as DNAs, coding and non-coding RNAs, lipids, and proteins that may have immunosuppressive effects or favor tumor invasion [6]. Among the non-coding RNAs, we have miRNAs, short RNAs of 18 to 22 nucleotides, which play a significant role in regulating gene expression at the post-transcriptional level [7].

This study presents the characterization of miRNAs contained in exosomes isolated from the plasma of a group of patients with gastric cancer and benign gastric pathologies. We reported a molecular signature of 9 miRNAs: of which, miR-92a-3p, miR451a, and miR126-3p were ranked as the most relevant in this study. Additionally, the target genes of the miRNAs found are described to associate them with biological processes and metabolic pathways involved in tumor development.

## Materials and methods

### Patient sample collection

The present study included 20 patients with primary gastric cancer at different stages who attended the Oncological Gastroenterology Unit of the Instituto Nacional de Cancerología, Bogotá, Colombia, and 10 controls with benign gastric pathologies confirmed by endoscopy biopsy no older than one year, who attended gastroenterology consultation at the Hospital Universitario de la Samaritana and the Centro Policlínico del Olaya in Bogotá, Colombia. The exclusion criteria for the study were having received any previous treatment, having previously had a neoplastic disease, having an active viral infection or autoimmune disease, having received a transplant, or being pregnant. Clinicopathological information was obtained by reviewing medical records.

After signing the informed consent form, 8 ml of peripheral blood was collected in tubes with EDTA as an anticoagulant to avoid platelet activation. The blood was centrifuged at 500g for 10 minutes to obtain the plasma. The ethics committee of each institution approved the study protocol in accordance with the Helsinki protocol. The characteristics of the study participants are summarized in Table 1.

**Table 1.**
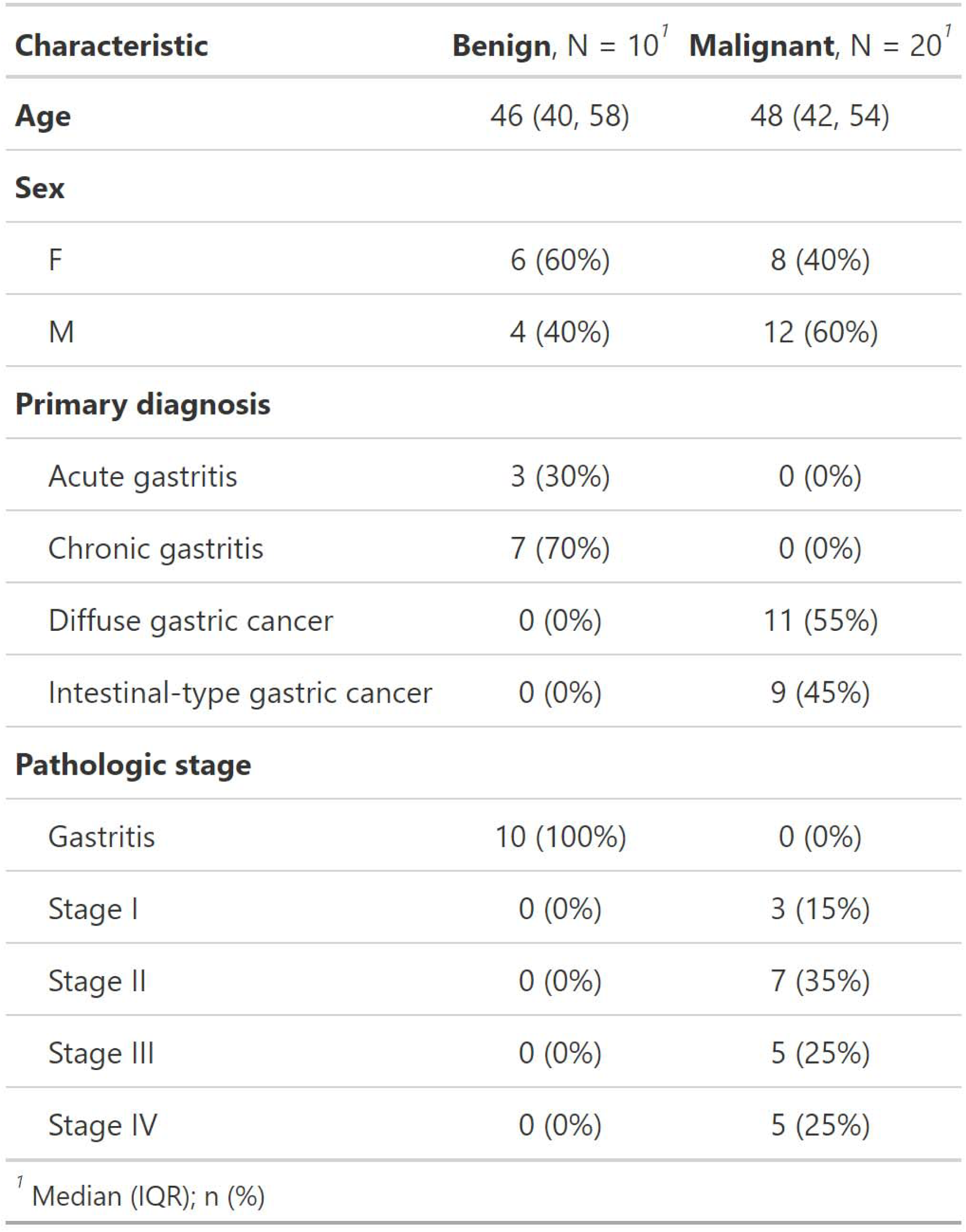
Clinical characteristics of study patients

### Extraction of EVs from human plasma

Exosomes were isolated from the plasma of patients with gastric cancer and individuals with a benign gastric disease by molecular exclusion chromatography, following the protocols of published studies with some modifications [8,9]. Plasma was thawed and centrifuged at 12,000g for 10 minutes; subsequently, the supernatant was passed through a 0.22 μm filter to remove contaminants such as bacteria and plasma proteins. For this study, qEV2 size exclusion columns were used (Izon Science, Christchurch, New Zealand). The columns were hydrated and cleansed with 1x phosphate buffer saline (PBS), sterile, and filtered through 0.22 μm. Once the column was stabilized, 2 ml of the filtered plasma was loaded onto it; 7 fractions of 2 ml were collected in Eppendorf tubes with low protein retention to avoid the loss of exosomes due to adhesion to the walls of traditional tubes.

### Confirmation of exosome collection

To confirm that what was obtained by molecular exclusion chromatography were exosomes, three methodologies were used: nanoparticle tracking analysis (NTA) to determine size and concentration, transmission electron microscopy (TEM) to identify the morphology of exosomes, and dot-blot assays to identify proteins on the exosome surface.

### Nanoparticle tracking analysis (NTA)

To evaluate particle size and concentration in the samples, the nanoparticle tracking analysis (NTA) was used, as described by Menezes-Neto [10]. This technique tracks particles based on light scattering and Brownian motion, determining the size of particles in a liquid suspension [11]. This study used the NanoSight LM10 system (Malvern Technologies, Malvern, UK), including a 638 nm laser, a CCD camera, and NTA software version 3.1 (build 3.1.46).

Samples were diluted 1:100 in 1X filtered PBS to reduce particle circulation in the capture fields and filtered PBS target was used as a blank. Three 1-minute videos were captured per sample.

### Transmission electron microscopy

To verify the presence of EVs in the fractions, 5 μl of fresh exosome concentrate was collected, put on Formvar-carbon-coated grids, and allowed to dry at room temperature for 5 minutes. Subsequently, negative staining was performed with 2% (v/v) of uranyl acetate in doubly distilled water for 5 minutes in the dark. Once the samples were dried, reads were made at 80 kV using the JEM-1400 Flash Electron Microscope equipped with a high-sensitivity sCMOS camera.

### Dot-blot assays for EV

Dot-blot assays were designed to detect membrane tetraspanins CD63 and CD81. Briefly, 5 μl of vesicles concentrate were added to 0.2 μm nitrocellulose membranes (Bio-Rad, 1620146) and allowed to dry at room temperature. The nonspecific sites were blocked in a standard TBS-T buffer supplemented with 5% non-fat milk for one hour. Subsequently, primary antibodies CD81 (Biolegend, 349504) and CD63 (Biolegend, 353008) were added and incubated at 4° C for 60 minutes. After washing the membranes with TBS-T buffer, they were incubated with a secondary Anti-IgG (HRP) antibody (ABCAM, ab6789) for 1 hour and washed again with TBS-T. Finally, Clarity Max Western ECL Substrate (Bio-Rad, 1705062) was added, and protein expression was detected by chemiluminescence in the ChemiDoc imaging system (Bio-Rad).

### Total RNA extraction

Total RNA was obtained from the exosome concentrate using the miRNeasy Mini Kit (Qiagen, Hilden, Germany. Cat. No./ID: 217004), following the manufacturer’s instructions. The RNA obtained was resuspended in 30 μl of RNase-free water; 800 μl of Trizol (Invitrogen Life Technologies, Carlsbad, CA) was added to a 200 μl sample of exosome concentrate, gently mixed, and incubated for 5 minutes at room temperature (15°C-20°C). Subsequently, 200 μl of chloroform was added, shaken vigorously for 15 seconds, and centrifuged for 15 minutes at 12,000 g at 4°C. The aqueous phase was transferred to the silica columns of the kit. The column was washed with 70% ethanol, allowed to dry at room temperature for 15 to 20 minutes, hydrated with 30 μl of RNase-free water, and centrifuged at 12,000 g for 2 minutes. The concentration and integrity of RNA purified from exosomes were evaluated using the Nanodrop® ND-200 (Thermo Fisher Scientific, Inc.) and the RNA 6000 Pico Kit (Agilent Technologies, Santa Clara, USA), respectively.

### RNA-seq analysis

Library construction and sequencing of the material obtained from the exosomes were carried out by BGI Small RNA Sequencing Services (Hong Kong). With 15 μL of the RNA concentrate, libraries were built using DNBSEQ (BGI Hong Kong), following the manufacturer’s instructions. The quality of the libraries was evaluated with a DNA chip (TapeStation, Agilent Technologies, Santa Clara, CA, USA).

Raw files in FastQ format from BGI Small RNA Sequencing Services (Hong Kong) were processed as summarized below. Quality control was performed with FastQC v0.11.9 to remove low-quality reads. The 5’ and 3’ sequencing adapters were removed using the Trim Galore tool, which implements Cutadapt v2.8 (https://github.com/marcelm/cutadapt) [12]. The reads were mapped against the reference genome (GRCh38.p13), and a count table was constructed with the identified small RNAs using the Spliced Transcripts Alignment to a Reference (STAR) software, with improved speed, sensitivity, and alignment accuracy, capable of mapping full-length RNA sequences [13]. A quality control and filter procedure were carried out; analyses were performed to determine the differential expression of miRNAs with the limma package [14,15]. To study different disease stages represented in the group of patients, comparisons were made between patients vs. non-tumor controls and between patients in different clinical stages vs. patients with benign gastric disease; miRNAs were considered differentially expressed with logFC>1 and a False Discovery Rate (FDR) of 5%.

### miRNA scoring

Based on the inter-group differential expression found in the study and conserving the sense of expression (up-down) in different comparisons, a score was assigned to the miRNAs with the best results in differential expression and reported in databases such as miRCancer (http://mircancer.ecu.edu/) and OncomiR (http://www.oncomir.org/), to generate greater robustness to the results.

### miRNA target prediction

To describe the miRNA target genes, three databases with different algorithms were used: miRNet (https://www.mirnet.ca/, accessed 20 August 2022), an online tool that integrates 20 databases related to interactions between miRNAs and other molecules [16]; TargetScan (https://www.targetscan.org/), for gene prediction based on complementary sequences conserved between miRNAs and the 3’ untranslated region (3’UTR) of miRNAs [17]; and miRDB (http://mirdb.org/) that makes predictions based on Machine Learning (ML) algorithms [18].

### TCGA stomach adenocarcinoma analysis

The TCGA stomach adenocarcinoma dataset (STAD-TCGA) was used to evaluate differentially expressed genes in tumor tissue. Briefly, the TCGA biolinks library (version 2.18.0) was used to obtain the raw counts necessary for the analysis [19]. Once the raw data were downloaded, DeSeq2 was used to perform differential gene expression analysis [20].

### Functional enrichment analysis

To determine the biological functions and metabolic pathways involved in target gene function, GeneTrail 3 (https://genetrail3.bioinf.uni-sb.de/) was used, a tool that integrates gene annotation in the Gene Ontology (GO), KEGG: Kyoto Encyclopedia of Genes and Genomes, WikiPathways, and Reactome pathway databases [21]. Metabolic terms and pathways with >10 hits and an adjusted p-value <0.005 were considered significant.

### Statistical analysis

Statistical analyses were refined in the R statistical programming language (https://www.r-project.org/). Differential expression was based on a linear model with limma, raw counts were transformed with the voom function to logCPM, normalizing the library sizes [15]. Statistically significant values in the gene expression study were those with an adjusted p-value <0.05. The functional enrichment analysis was performed with Fisher’s exact test, and the p-value was adjusted using the Benjamini-Hochberg method.

For the multivariate analysis, the Multiple correspondence analysis (MCA) and Principal Components Analysis (PCA) of the FactoMineR package [22] were used for the visualization factoextra. Logistic regression for benign and cancer patients was performed on the coordinates of the first two principal components of the PCA with the glm function, with a binomial response. Receiver operating characteristic (ROC) curves were used to assess the specificity and sensitivity of the model obtained with the pROC package [23]. The regression model was validated by LOOCV, using AUC as the main statistic.

For comparison of these two groups of patients, individual coordinates on the first principal component were used to perform a Wilcoxon exact test with the function wilcox.test, after verifying the assumptions of normality and equality of variances, using, respectively, Shapiro Wilk and the Levene test, the latter from the car package.

## Results

### Characterization of plasma-derived EVs

For the characterization of EVs, the recommendations of the International Society for Extracellular Vesicles (ISEV) were followed to perform studies on EVs (MISEV)[24]. By nanoparticle tracking analysis (NTA), a concentration of 5.84×109 particles/ml was determined with an average size of less than 200 nm in diameter (Figure 1). Transmission electron microscopy (TEM) revealed membrane-surrounded structures with a diameter between 30 and 150 nm. The expression of CD81 and CD63 was evidenced by dot-blot assay and flow cytometry in plasma-derived exosomes from gastric cancer patients.

**Figure 1.**
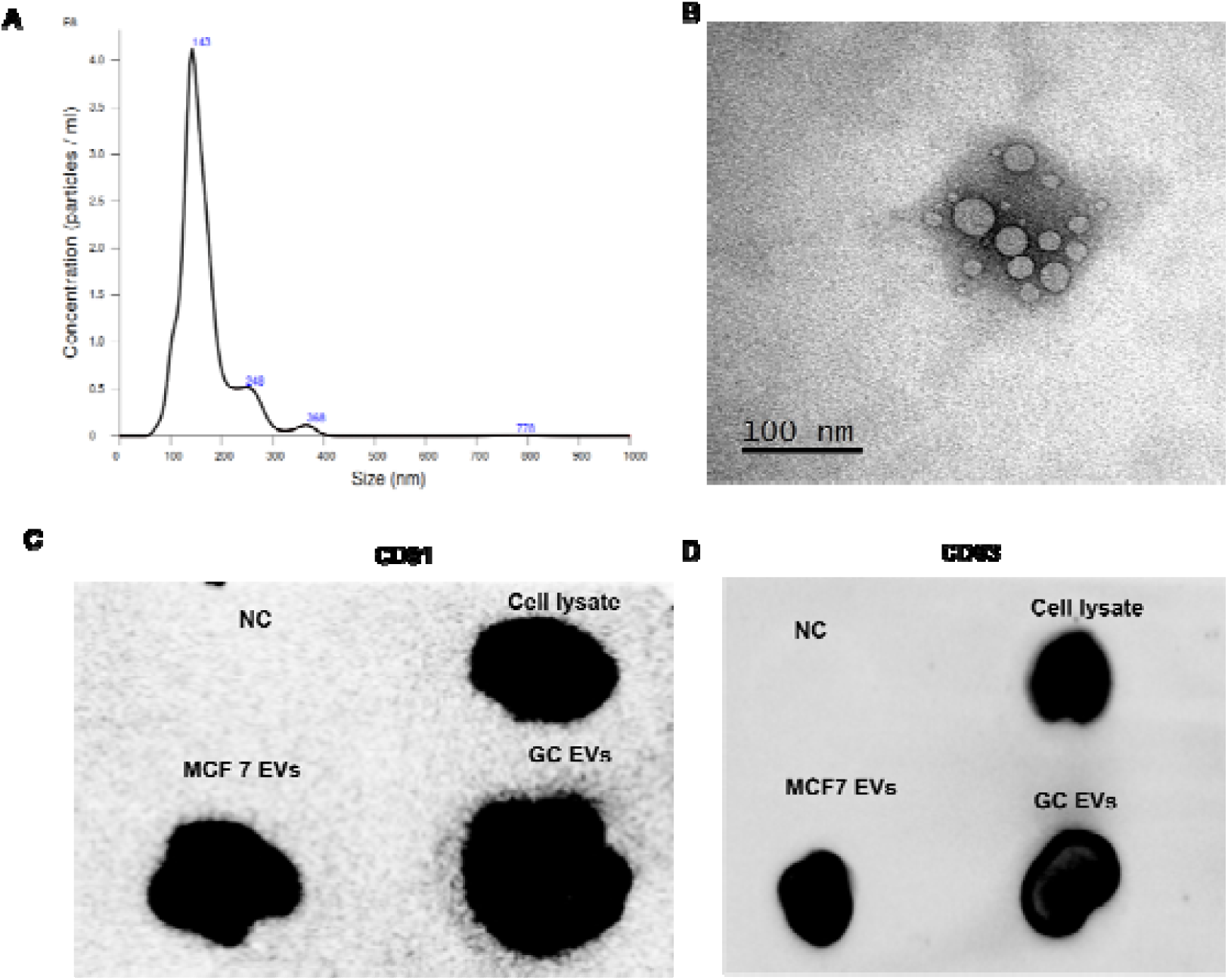
This study employs several techniques to investigate the properties of exosomes, including: **(A)** NanoTracking Analysis, which allows for the estimation of particle number and size; **(B)** Transmission Electron Microscopy, which enables the evaluation of vesicular structures at high resolution; and **(C)** Dotblot, a method used to verify the presence of proteins associated (tetraspanins CD81 and CD63) with exosome membranes.

### Exosomal sRNA expression profiles

This study included exosome samples from the plasma of 20 patients with gastric cancer and 10 controls with benign gastric pathology. Small RNA-seq was performed, and an average of 1.28 million unique reads per sample were obtained. Approximately 32.2% of the reads were mapped to the genome. On average, 156 mature miRNAs were found in exosome fractions from gastric cancer patients and controls (Figure 2), and those used for differential expression studies were selected. Raw counts were calculated as CPM (count per million).

**Figure 2.**
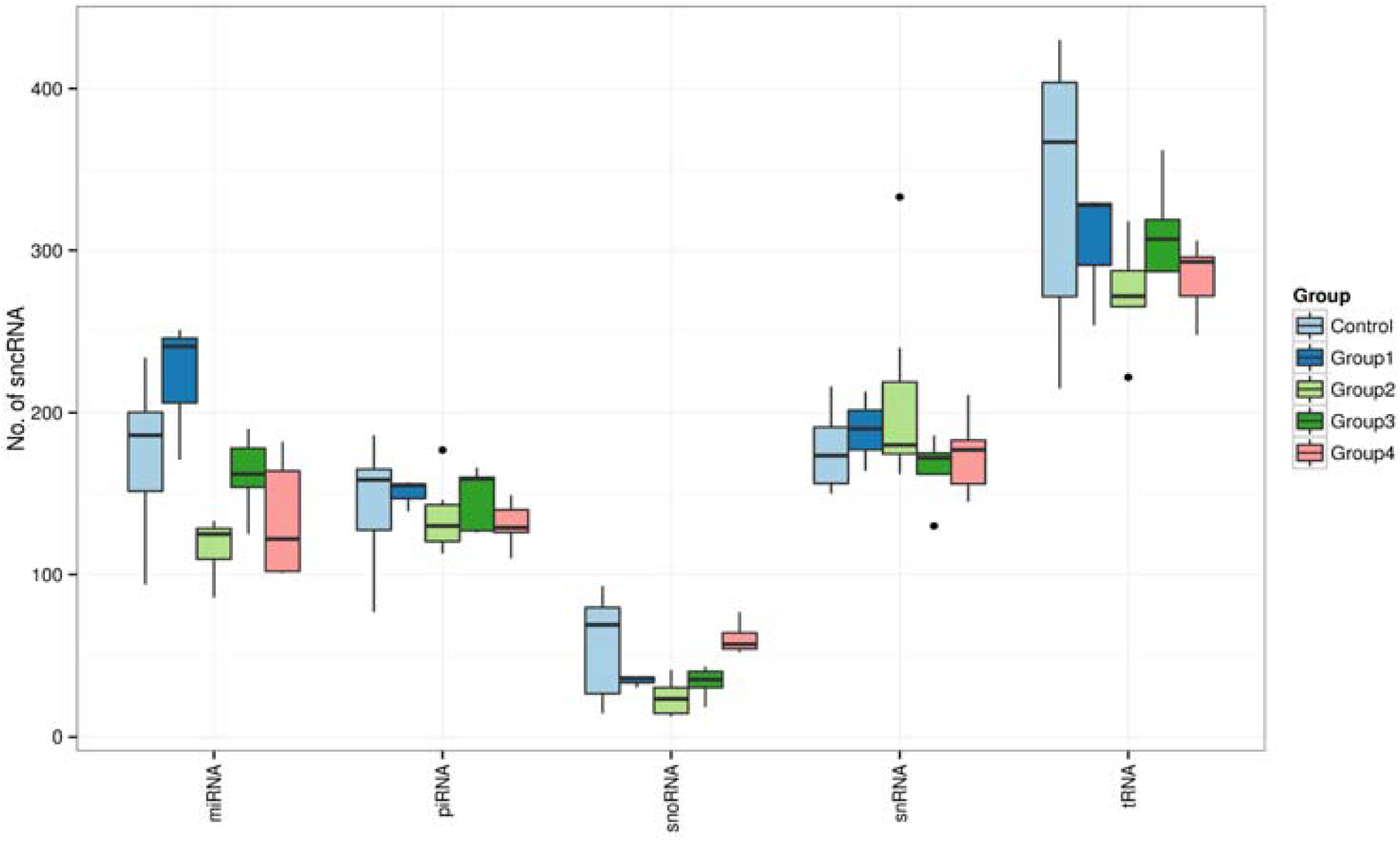
This boxplot shows the distribution of unique small non-coding RNA (sncRNA) between different patient groups. The patient groups include four groups with gastric cancer (group 1-4), as well as a control group with benign gastric diseases.

To determine which miRNAs were differentially expressed, the tally table was filtered, and miRNAs with less than 10 counts per sample were eliminated and samples with atypical data were deleted from the final analysis. Initially, we evaluated the overall differential expression between patients diagnosed with cancer and those diagnosed with benign gastric disease. As a result, we identified a molecular signature of 9 miRNAs that were differentially expressed. Additionally, we conducted a complementary analysis by comparing subgroups of patients with gastric cancer at different stages of the disease against subjects with benign disease. This analysis revealed 16 miRNAs that were differentially expressed, with 13 being down-regulated and 2 being up-regulated. (Table S1).

Of these 15 miRNAs, three (hsa-miR-92a-3p, hsa-miR-451a, and hsa-miR-126-3p) were ranked as the best candidates, we call them winners according to our classification (see methods section), to be evaluated in a representative cohort to determine their sensitivity and specificity and propose them as biomarkers for gastric cancer (Figure 3). The winning miRNAs were chosen to perform a search for target genes and subsequent functional enrichment.

**Figure 3.**
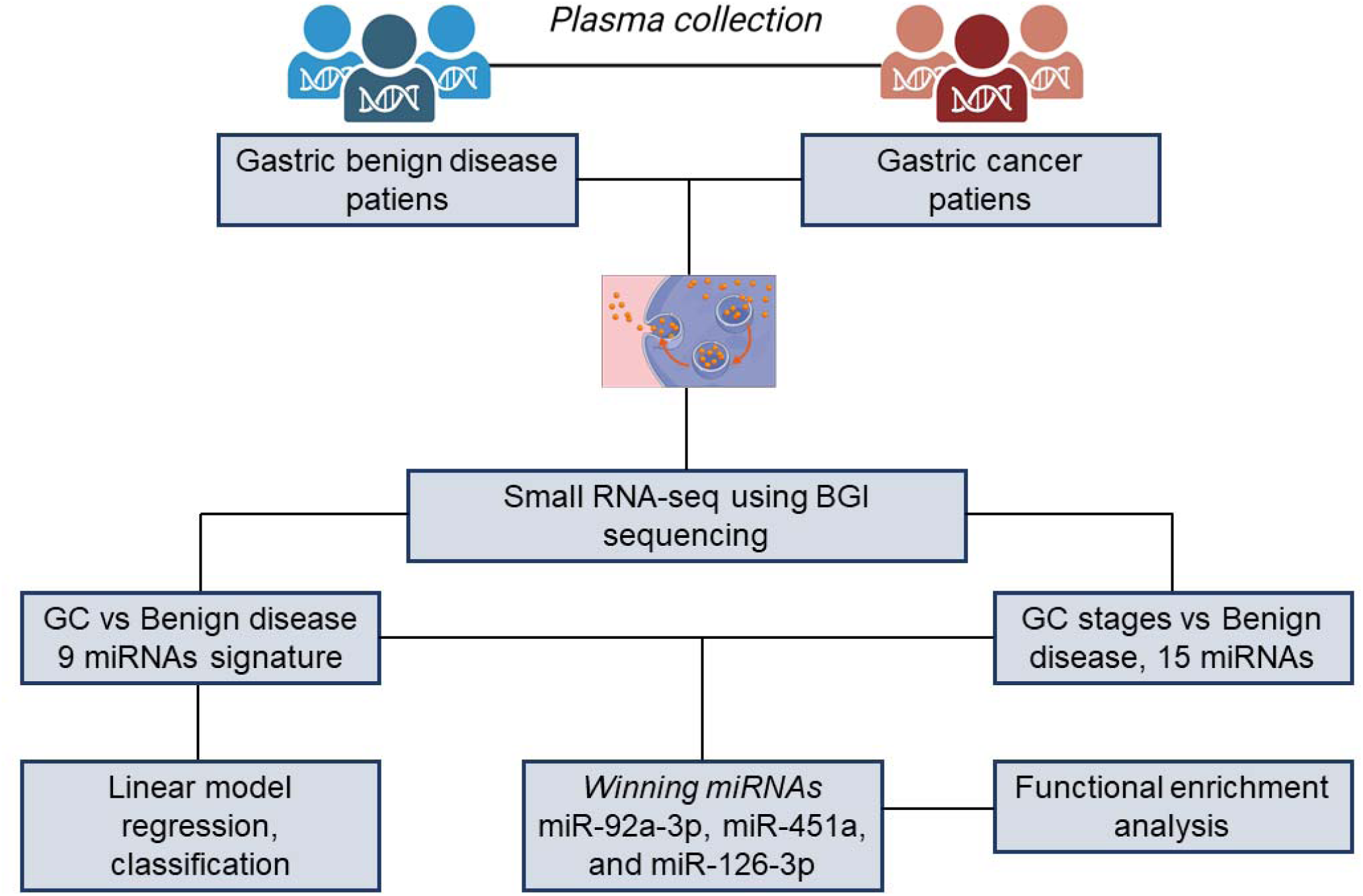
Pipeline used to analyze miRNAs derived from EVs in two patient groups: benign gastric disease and gastric cancer. EVs were isolated, and small RNA-seq was performed to quantify miRNAs present in both populations. We discovered a differential signature of 9 miRNAs, and by comparing different stages of cancer, we identified 15 miRNAs with differential expression. After scoring we propose three winning miRNAs as potential biomarkers of gastric cancer.

### miRNA target genes

The winning miRNAs were used to search in miRNet, TargetScan, and miRDB target genes. Below is a summary of the predictions of the applied algorithms (Supplementary Table S2).

### Validation in TCGA cohort Exosomal miRNA targets

After differential gene expression was applied with DESeq2, with a threshold of logFC >2 and adjusted p-value <0.01, we found 454 dysregulated genes and 469 over-expressed genes (Figure 4A). This analysis aimed to identify which miRNA target genes of our cohort are altered in the TCGA reference dataset. In this sense, of the 1,618 potential miRNA target genes, 42 genes were found in our TCGA analysis results (Supplementary Table S3).

**Figure 4.**
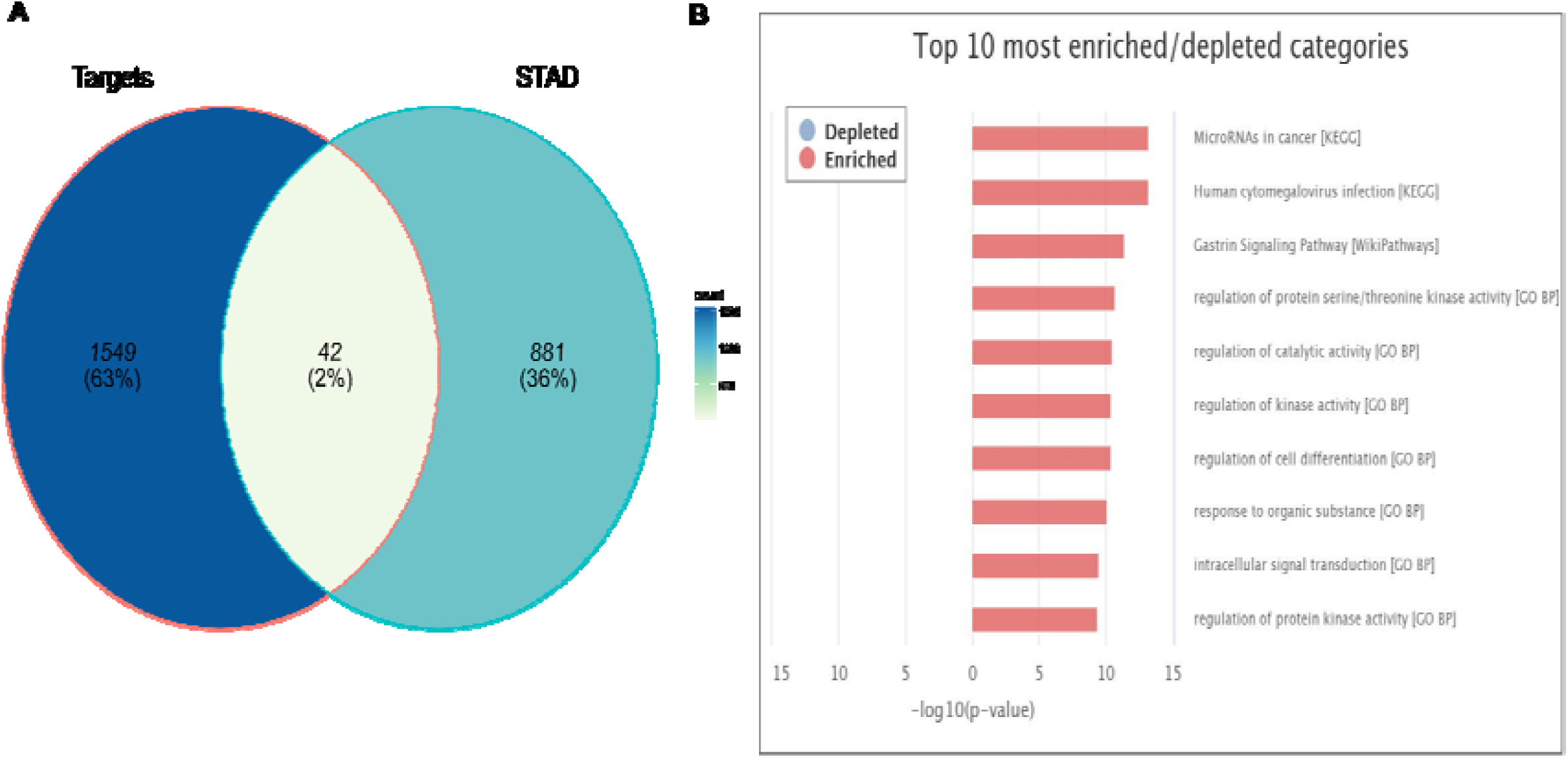
Venn diagram of the analysis of TCGA and winning miRNA targets and functional enrichment of winning miRNA targets. **(A)** miRNA target genes derived from gastric cancer EVs and their presence in the differentially expressed genes of STAD-TCGA. **(B)** Top 10 biological processes (GO-BP) and metabolic pathways (KEGG-Wikipathways) represented in the target genes of hsa-miR-451a, hsa-miR-126-3p, and hsa-miR-92a-3p

### Functional enrichment

After identifying the potential target genes of hsa-miR-451a, hsa-miR-126-3p, and hsa-miR-92a-3p, the highest-ranking metabolic pathway was miRNAs in cancer (KEGG); interestingly, within the top metabolic pathways represented, we found the gastrin signaling pathway (Wikipathways), a pathway with special interest in the development of gastric cancer and other gastrointestinal tumors (Figure 4B). We also found other biological processes (GO-BP) associated with kinase-associated cell signaling and cell differentiation (Supplementary file 2).

### Multivariate Analysis

To relate the clinical variables to the transcriptomic results, a multiple correspondence analysis (MCA) was performed, to determine that the study subjects were separated based on the type of disease (benign-malignant) (Figure 5A). To find clusters based on clinical variables, the k-means clustering method was used. On the other hand, with the 9 miRNAs signature with differential expression in the study groups, a PCA was done and it was possible to determine that the miRNAs have a high correlation between them and, in turn, with the presence of the outcome (benign gastric disease or gastric cancer), complementing the findings of the clinical variables in the MCA (Figure 5B).

**Figure 5.**
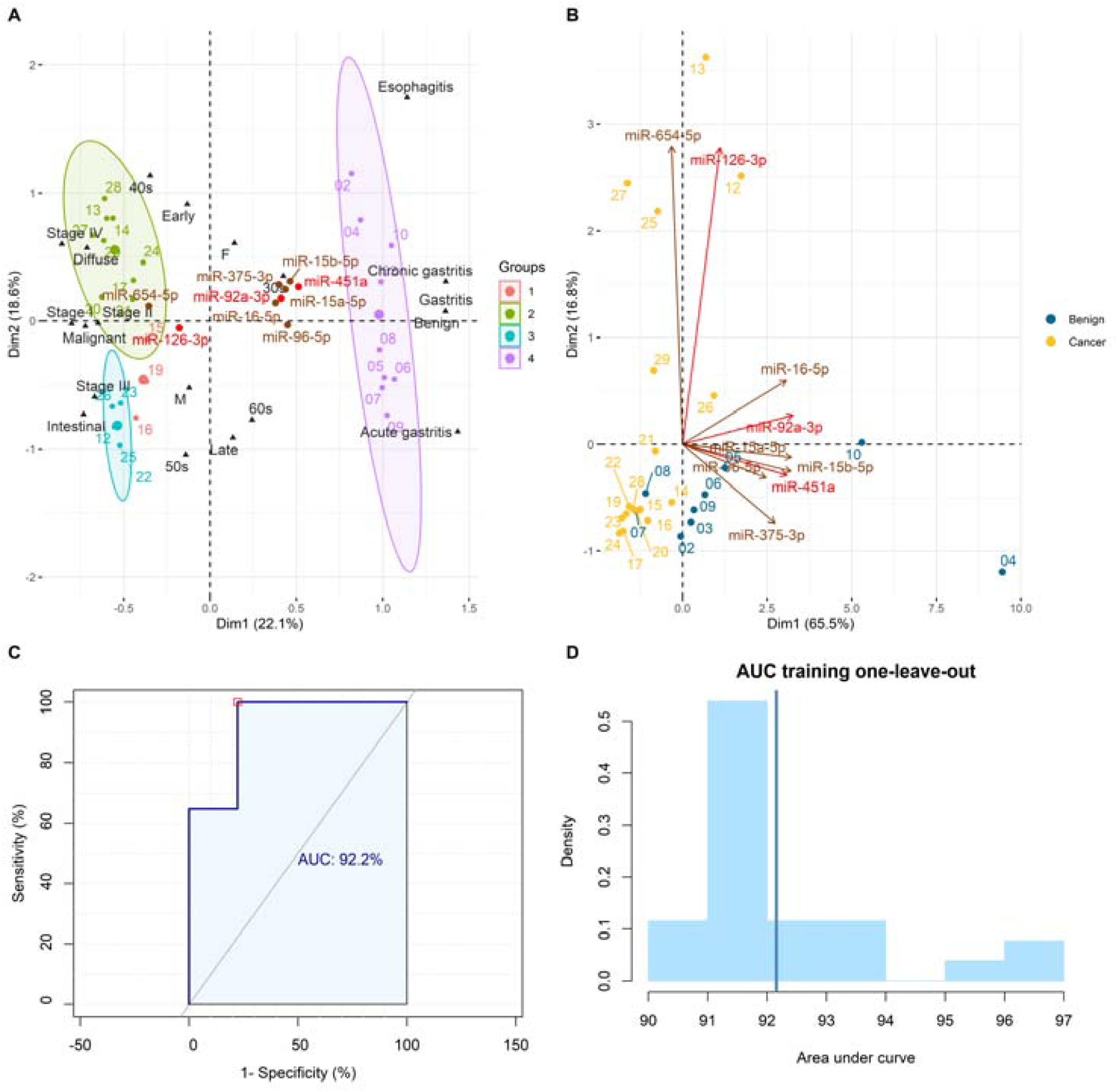
Multivariate analysis and classification model based on logistic regression. **(A)** Clustering of the clinicopathological variables of the subjects. **(B)** Reduction of the dimensionality of the molecular signature of miRNAs in EVs and its correlation with the type of disease (Cancer-Benign disease). **(C)** ROC curve with the molecular signature as testing set. **(D)** LOOCV analysis for the performance of the model with the training set.

### Logistic Regression and Cross-Validation

Using the coordinates of the individuals in the first two components of the PCA, logistic regression was implemented to determine if the summary variables have sufficient power to classify between individuals with the benign gastric disease and gastric cancer. The logistic regression for dimension 1, yielded a significance of 0.0147*, explaining that the molecular signature of 9 miRNAs can discriminate patients with the two types of disease. To confirm this result, the Wilcoxon rank-sum exact test was performed (after verification of homoscedasticity) and a significant difference was obtained between the two conditions (p-value 0.0131).

To evaluate the prediction of the set of miRNAs with differential expression (predictors) to classify all patients in the two types of disease (response), an analysis of ROC curves was used, taking the statistical AUC to measure the prediction of the quality. (AUC=92.16%, Figure 5C). In order to evaluate the generality of the model and rule out extreme influences from any of the patients, the leave-one-out cross-validation (LOOCV) method was used, obtaining AUC values between 90%-97% (Figure 5D).

## Discussion

Cellular communication mediated by EVs plays a significant role in physiological and pathological processes. EVs can be formed from the cell membrane and undergo a maturation process through the vesicular transport system [25]. Cellular communication mediated by EVs in the microenvironment or over long distances is a mechanism used by tumor cells to modify cell behavior and induce molecular changes in adjacent epithelial cells, immune system cells, and endothelial cells, among others [26].

In this exploratory study, we characterized the exosomes associated miRNAs in patients with gastric cancer, using an adjusted methodology to obtain EVs with new-generation technologies. miRNAs can modulate gene expression in target cells at a post-transcriptional level, modulating physiological processes in the recipient cell [27]. Our study is particularly relevant because gastric cancer is a disease that lacks biomarkers for early diagnosis.

The methodology applied, isolation of EVs by SEC from human plasma, proved to be adequate for the characterization of these structures by NTA and TEM, evidencing an approximate size of 150 nm. The presence of cell membrane tetraspanins CD81 and CD63, recognized as typical of EVS, was confirmed by dot-blot assays. Finally, sufficient RNA was obtained to perform small RNA-seq studies.

Here we present a molecular signature composed of 9 miRNAs with a potential classification power between patients with gastric cancer and patients with the benign gastric disease (ROC-AUC 92.2%). To validate these results, a LOOCV was applied, in which an optimum AUC of 90%-97% was maintained. It is necessary to use a cohort of different patients to corroborate the classification capacity between the two types of pathologies, which can constitute a significant advance in the achievement of biomarkers for gastric cancer.

We describe the differential expression of three winner miRNAs, two with over-expression (hsa-miR- 92a-3p, hsa-miR-451a) and one dysregulated (hsa-miR-126-3p), compared to sex- and age-matched non-tumor controls. In the molecular signature found in this study, hsa-miR-92a-3p has been described as a promoter of cell proliferation in breast cancer cells by modulating Krüppel-like transcription factors (KLFs) [28]. Regarding the inclusion of this miRNA in EVs, a recent study by Uehara et al. uses a mouse model to demonstrate that miRNA-92a-3p contained in EVs promotes osteolytic metastasis by directly blocking PTEN in mature osteoclasts [29]. As a potential biomarker in gastric cancer, hsa-miR-92a-3p was found to be overexpressed in plasma in a cohort of 160 patients with gastric cancer and validated in independent cohorts as an early diagnostic marker.

In this same study, which identified a molecular signature of five miRNAs, hsa-miR-451a was also found to be over-expressed in gastric cancer patients [30]. In EVs, hsa-miR-451a was described by Yoshida et al. in bile-derived exosomes from patients with biliary tract cancers (BTCs) and functioning as a potential biomarker of these diseases [31]. Like miRNA-92a-3p, hsa-miR-451a also targets and suppresses the expression of the PTEN tumor suppressor in neuroendocrine tumors of the rectum, generating invasion and metastasis [32].

On the other hand, hsa-miR-126-3p, a miRNA described as a tumor suppressor dysregulated in various types of cancer, in certain cases, due to the methylation of the CpG islands of its promoter [33,34]. In addition, platelet-derived EVs contain hsa-miR-126-3p with a blocking activity of the PI3K/AKT pathway in breast tumor cells, which is an attractive strategy for cancer therapy [35]. These findings were also validated in the study by Di Paolo et al., who, in a xenographic model of lung cancer, using an hsa-miR-126-3p mimic encapsulated in lipid microspheres, achieved a decrease in tumor proliferation through blockade in the PI3KR2-AKT pathway [36].

Regarding the biological processes and metabolic pathways potentially affected by the miRNA target genes described here, the metabolic pathway of miRNAs in cancer (hsa05206) is represented with 22 genes. This metabolic pathway is crucial in the post-transcriptional regulation of tumor suppressor genes and oncogenes, favoring proliferation, cell differentiation, and apoptosis [37]. Another metabolic pathway with particular representation in the target genes was the gastrin signaling pathway, a hormone produced by G cells in the antropyloric portion of the stomach, with an important role in gastric acid secretion and gastric epithelial cell proliferation [38]. Gastrin has also been associated with the development of gastric cancer, primarily as a growth factor that activates various pathways mediated by the epidermal growth factor receptor (EGFR) and PI3K and MAPK kinases [39,40].

In the search for biomarkers that optimize the diagnosis, prognosis, and follow-up of gastric cancer, a tool of recent interest is liquid biopsy, which is based on seeking information on tumor biology [41]. The most studied sources of information in liquid biopsy are CTCs, cell free DNA (cfDNA), cytokines, and EVs. Due to their lipid bilayer, EVs are important for nucleic acid transport, in particular miRNAs protecting them from enzymatic degradation in the circulation [42]. In gastric cancer, the most important sources for EV collection are peripheral blood, peritoneal fluid, ascites, and gastric juice [43].

Our study present the characterization of the exosomes contained miRNAs from patients with gastric cancer. This is a preliminary study with a small number of patients and a study with a larger cohort of patients must be performed allowing the validation of the molecular signature found here, and providing molecules to be postulated with greater certainty as candidates for disease biomarkers.

## Conclusions

This study of EVs in gastric cancer patients with different disease stages provided relevant information on the molecular signature carried by this type of vesicles in blood circulation. The hsa-miR-92a-3p, hsa-miR-126-3p and hsa-miR-451a miRNAs were studied as molecules with potential effects on the development of cancer. Some of these biological functions were already reported in other studies, which denotes the importance of these molecules in disease development and progression. We hope that these results can be validated in a study with a larger number of patients and contribute to the discovery of biomarkers in liquid biopsy, a necessary tool for the control of gastric cancer.

## Funding

This research was funded by Instituto Nacional de Cancerología (Grant Number XRPM:C19010300- 454 to J.A.R.), Colombian Sciences Ministry scholarship (BB-2019-01 to A.R.R.).

**Conflict of Interest Statement:** None declared

## Data availability

The RNA sequencing expression data underling this article have been deposited in the National Center for Biotechnology Information’s (NCBIs) Gene Expression Omnibus (GSE227778).

## Abbreviations

TCGA: The Cancer Genome Atlas
LOOCV: Leave-One-Out Cross-Validation
AUC: Area Under the Curve
MISEV: Minimal Information for Studies of Extracellular Vesicles
nm: Nanometers
sRNA: Small RNA
STAD: Stomach adenocarcinoma
ROC: Receiver Operating Characteristic
PTEN: Phosphatase and Tensin homolog
PI3K: Phosphoinositide 3-kinase
AKT: serine/threonine kinase
MAPK: Mitogen-activated protein kinase

## Notes

### Competing Interest Statement

The authors have declared no competing interest.

