## Supplementary file 1 for "Exosomal microRNA signature from plasma-derived extracellular vesicles in gastric cancer"

**Table S1.** Importance ranking of miRNAs that showed differential expression (logFC>2 & adjusted p-value <0.05), using the scoring method described above. This ranking allowed us to identify the most relevant miRNAs in our study and prioritize them as winners for further analysis.

| **miRNA** | **Sense expression** | **Adjusted p-value** | **Score** |
| --- | --- | --- | --- |
| **hsa-miR-92a-3p** | **Down** | **<0.001** | **5** |
| **hsa-miR-451a** | **Down** | **<0.001** | **4** |
| **hsa-miR-126-3p** | **Up** | **<0.001** | **3** |
| hsa-miR-375-3p | Down | <0.001 | 1 |
| hsa-miR-16-5p | Down | <0.001 | 1 |
| hsa-miR-654-5p | Up | <0.001 | 1 |
| hsa-miR-15b-5p | Down | <0.001 | 0 |
| hsa-miR-96-5p | Down | <0.001 | 3 |
| hsa-miR-15a-5p | Down | <0.001 | 0 |
| hsa-miR-138-5p | Down | <0.001 | 2 |
| hsa-miR-345-5p | Down | 0.001 | 1 |
| hsa-miR-424-3p | Down | 0.01 | 0 |
| hsa-miR-3173-5p | Down | 0.03 | 1 |
| hsa-let-7a-2-3p | Down | 0.02 | 2 |
| hsa-miR-29b-3p | Down | 0.04 | 0 |

**Table S2.** Distribution of potential targets for winning miRNAs derived from patients with gastric cancer

| **miRNA** | **Total gene targets** |
| --- | --- |
| hsa-miR-92a-3p | 1,482 |
| hsa-miR-451a | 66 |
| hsa-miR-126-3p | 70 |

**Table S3.** Exosomal-miRNA target genes with differential expression in the STAD-TCGA cohort

| **Gene** | **logFC** | **Gene** | **logFC** |
| --- | --- | --- | --- |
| NEK2 | 2.95 | SOX2 | -3.58 |
| GNG7 | -2.65 | CHST9 | -3.31 |
| KIFC1 | 2.24 | MUC21 | -3.21 |
| EPHA7 | -2.63 | CIDEC | -2.38 |
| HOXC8 | 4.66 | BMP8A | 2.15 |
| AURKA | 2.1 | OXTR | 2.26 |
| RCC1 | 2.94 | DTL | 2.3 |
| CDC6 | 2.72 | KIF2C | 2.32 |
| GPX3 | -2.67 | CBX2 | 2.42 |
| CDC20 | 2.53 | NCAPH | 2.47 |
| SOX11 | 2.55 | MMP9 | 2.5 |
| CDK1 | 2.39 | POLQ | 2.73 |
| KIF1A | -2.49 | KIF20A | 2.81 |
| HAND2 | -2.34 | AURKB | 2.85 |
| MKI67 | 2.95 | HOXA9 | 2.97 |
| NOX4 | 2.77 | HMGA2 | 3.07 |
| CKB | -2.48 | ZNF695 | 3.1 |
| EPHB2 | 2.95 | CLSPN | 3.2 |
| HBB | -2.91 | ZIC5 | 3.65 |
| HOXA13 | 6.15 | KIF18B | 3.66 |
| MYBL2 | 3.36 | MMP7 | 3.95 |
