## Supplementary file 2 for "Exosomal microRNA signature from plasma-derived extracellular vesicles in gastric cancer"

### geneList.ora.zip

#### Reactome_-_Pathways

| Type | Name | #Hits | Expected score | p-Value |
|---|---|---|---|---|
| enriched | Interleukin-4 and Interleukin-13 signaling | 11 | 1.465720 | 0.000055 |
| enriched | Cyclin E associated events during G1/S transition | 6 | 0.380001 | 0.000190 |
| enriched | Synthesis of PIPs at the plasma membrane | 7 | 0.719288 | 0.000518 |
| enriched | Estrogen-dependent nuclear events downstream of ESR-membrane signaling | 5 | 0.325715 | 0.000836 |
| enriched | Cyclin A:Cdk2-associated events at S phase entry | 5 | 0.407144 | 0.002097 |
| enriched | FOXO-mediated transcription of cell cycle genes | 4 | 0.230715 | 0.002413 |
| enriched | Oncogene Induced Senescence | 5 | 0.447858 | 0.002413 |

#### KEGG_-_Pathways

| Type | Name | #Hits | Expected score | p-Value |
|---|---|---|---|---|
| enriched | Human cytomegalovirus infection | 25 | 3.053580 | 0.000000 |
| enriched | MicroRNAs in cancer | 22 | 2.185010 | 0.000000 |
| enriched | PI3K-Akt signaling pathway | 25 | 4.790730 | 0.000000 |
| enriched | FoxO signaling pathway | 16 | 1.777860 | 0.000000 |
| enriched | Small cell lung cancer | 14 | 1.248570 | 0.000000 |
| enriched | Pathways in cancer | 30 | 7.179300 | 0.000000 |
| enriched | Hepatitis B | 17 | 2.198580 | 0.000000 |
| enriched | Chronic myeloid leukemia | 12 | 1.031430 | 0.000000 |
| enriched | Prostate cancer | 13 | 1.316430 | 0.000000 |
| enriched | Cellular senescence | 15 | 2.171430 | 0.000000 |
| enriched | Growth hormone synthesis, secretion and action | 13 | 1.615000 | 0.000000 |
| enriched | Human T-cell leukemia virus 1 infection | 17 | 2.972150 | 0.000000 |
| enriched | EGFR tyrosine kinase inhibitor resistance | 11 | 1.072150 | 0.000000 |
| enriched | Neurotrophin signaling pathway | 12 | 1.615000 | 0.000001 |
| enriched | AGE-RAGE signaling pathway in diabetic complications | 11 | 1.357150 | 0.000001 |
| enriched | mTOR signaling pathway | 13 | 2.076430 | 0.000002 |
| enriched | Kaposi sarcoma-associated herpesvirus infection | 14 | 2.524290 | 0.000003 |
| enriched | Chemokine signaling pathway | 14 | 2.565010 | 0.000003 |
| enriched | TNF signaling pathway | 11 | 1.520000 | 0.000004 |
| enriched | Estrogen signaling pathway | 12 | 1.872860 | 0.000004 |
| enriched | Epstein-Barr virus infection | 14 | 2.714290 | 0.000006 |
| enriched | Yersinia infection | 11 | 1.628580 | 0.000006 |
| enriched | Gastric cancer | 12 | 1.995010 | 0.000006 |
| enriched | Endometrial cancer | 8 | 0.787145 | 0.000009 |
| enriched | Bladder cancer | 7 | 0.556430 | 0.000009 |
| enriched | Relaxin signaling pathway | 11 | 1.750720 | 0.000011 |
| enriched | Parathyroid hormone synthesis, secretion and action | 10 | 1.438580 | 0.000012 |
| enriched | ErbB signaling pathway | 9 | 1.153570 | 0.000015 |
| enriched | Fluid shear stress and atherosclerosis | 11 | 1.872860 | 0.000019 |
| enriched | Longevity regulating pathway | 9 | 1.207860 | 0.000021 |

| | | | | |
|---|---|---|---|---|
| enriched | Viral carcinogenesis | 13 | 2.727860 | 0.000023 |
| enriched | Sphingolipid signaling pathway | 10 | 1.615000 | 0.000030 |
| enriched | Human immunodeficiency virus 1 infection | 13 | 2.863580 | 0.000037 |
| enriched | Endocrine resistance | 9 | 1.330000 | 0.000041 |
| enriched | Pancreatic cancer | 8 | 1.017860 | 0.000042 |
| enriched | Hepatitis C | 11 | 2.103580 | 0.000046 |
| enriched | Human papillomavirus infection | 16 | 4.465010 | 0.000060 |
| enriched | Apoptosis | 10 | 1.832150 | 0.000078 |
| enriched | cGMP-PKG signaling pathway | 11 | 2.266430 | 0.000087 |
| enriched | Proteoglycans in cancer | 12 | 2.727860 | 0.000095 |
| enriched | Colorectal cancer | 8 | 1.167150 | 0.000100 |
| enriched | Cholinergic synapse | 9 | 1.533580 | 0.000105 |
| enriched | GnRH secretion | 7 | 0.868574 | 0.000108 |
| enriched | Non-small cell lung cancer | 7 | 0.895717 | 0.000130 |
| enriched | Breast cancer | 10 | 1.995010 | 0.000137 |
| enriched | Regulation of actin cytoskeleton | 12 | 2.890720 | 0.000146 |
| enriched | Central carbon metabolism in cancer | 7 | 0.936431 | 0.000159 |
| enriched | Renal cell carcinoma | 7 | 0.936431 | 0.000159 |
| enriched | Melanoma | 7 | 0.977145 | 0.000206 |
| enriched | Glioma | 7 | 1.017860 | 0.000262 |
| enriched | Chagas disease (American trypanosomiasis) | 8 | 1.384290 | 0.000274 |
| enriched | C-type lectin receptor signaling pathway | 8 | 1.411430 | 0.000309 |
| enriched | Focal adhesion | 11 | 2.700720 | 0.000317 |
| enriched | Hepatocellular carcinoma | 10 | 2.266430 | 0.000335 |
| enriched | Autophagy - animal | 9 | 1.859290 | 0.000356 |
| enriched | Insulin signaling pathway | 9 | 1.859290 | 0.000356 |
| enriched | Insulin resistance | 8 | 1.465720 | 0.000364 |
| enriched | Measles | 9 | 1.872860 | 0.000364 |
| enriched | HIF-1 signaling pathway | 8 | 1.479290 | 0.000379 |
| enriched | Leukocyte transendothelial migration | 8 | 1.520000 | 0.000450 |
| enriched | PD-L1 expression and PD-1 checkpoint pathway in cancer | 7 | 1.221430 | 0.000681 |
| enriched | AMPK signaling pathway | 8 | 1.628580 | 0.000701 |
| enriched | Acute myeloid leukemia | 6 | 0.909288 | 0.000802 |
| enriched | Cell cycle | 8 | 1.682860 | 0.000802 |
| enriched | Cushing syndrome | 9 | 2.117150 | 0.000802 |
| enriched | GnRH signaling pathway | 7 | 1.262150 | 0.000802 |
| enriched | Platelet activation | 8 | 1.682860 | 0.000802 |
| enriched | Shigellosis | 6 | 0.909288 | 0.000802 |
| enriched | Apelin signaling pathway | 8 | 1.859290 | 0.001522 |
| enriched | Toll-like receptor signaling pathway | 7 | 1.425000 | 0.001522 |
| enriched | Signaling pathways regulating pluripotency of stem cells | 8 | 1.900010 | 0.001726 |
| enriched | MAPK signaling pathway | 12 | 3.990010 | 0.001854 |
| enriched | B cell receptor signaling pathway | 6 | 1.085720 | 0.001943 |
| enriched | Toxoplasmosis | 7 | 1.520000 | 0.002115 |
| enriched | Adrenergic signaling in cardiomyocytes | 8 | 2.022150 | 0.002421 |
| enriched | Transcriptional misregulation in cancer | 9 | 2.510720 | 0.002421 |
| enriched | VEGF signaling pathway | 5 | 0.800716 | 0.002935 |
| enriched | Long-term depression | 5 | 0.827859 | 0.003368 |
| enriched | Aldosterone-regulated sodium reabsorption | 4 | 0.502144 | 0.003518 |
| enriched | Longevity regulating pathway - multiple species | 5 | 0.841431 | 0.003518 |
| enriched | Thyroid cancer | 4 | 0.502144 | 0.003518 |
| enriched | Pathogenic Escherichia coli infection | 9 | 2.727860 | 0.003955 |
| enriched | Progesterone-mediated oocyte maturation | 6 | 1.289290 | 0.004176 |
| enriched | Fc epsilon RI signaling pathway | 5 | 0.909288 | 0.004753 |

### GO_-_Cellular_Component

| Type | Name | #Hits | Expected score | p-Value |
|---|---|---|---|---|
| enriched | cytoplasm | 98 | 60.705200 | 0.000037 |
| enriched | cytosol | 105 | 68.427300 | 0.000069 |
| enriched | cell junction | 39 | 17.398600 | 0.000139 |
| enriched | cellular component | 262 | 242.115000 | 0.000139 |
| enriched | cell-substrate adherens junction | 17 | 5.428590 | 0.001108 |
| enriched | cell-substrate junction | 17 | 5.482870 | 0.001108 |
| enriched | focal adhesion | 17 | 5.387870 | 0.001108 |
| enriched | intracellular organelle | 162 | 129.160000 | 0.001108 |
| enriched | membrane-bounded organelle | 162 | 130.042000 | 0.001108 |
| enriched | nucleoplasm | 76 | 49.332300 | 0.001108 |
| enriched | postsynaptic density | 13 | 3.460720 | 0.001108 |
| enriched | postsynaptic specialization | 13 | 3.487870 | 0.001108 |
| enriched | nucleus | 97 | 68.291600 | 0.001150 |
| enriched | chromatin | 30 | 14.263600 | 0.001767 |
| enriched | early endosome | 12 | 3.270720 | 0.001947 |
| enriched | adherens junction | 19 | 7.233590 | 0.002038 |

### GO_-_Biological_Process

| Type | Name | #Hits | Expected score | p-Value |
|---|---|---|---|---|
| enriched | regulation of protein serine/threonine kinase activity | 34 | 6.812880 | 0.000000 |
| enriched | regulation of catalytic activity | 76 | 30.861500 | 0.000000 |
| enriched | regulation of cell differentiation | 65 | 24.170800 | 0.000000 |
| enriched | regulation of kinase activity | 43 | 11.522200 | 0.000000 |
| enriched | response to organic substance | 72 | 29.232900 | 0.000000 |
| enriched | intracellular signal transduction | 59 | 21.727900 | 0.000000 |
| enriched | regulation of protein kinase activity | 39 | 10.490700 | 0.000000 |
| enriched | positive regulation of cellular metabolic process | 91 | 44.256500 | 0.000000 |
| enriched | positive regulation of metabolic process | 96 | 48.287300 | 0.000000 |
| enriched | regulation of phosphorylation | 57 | 21.185100 | 0.000000 |
| enriched | regulation of signaling | 94 | 47.364400 | 0.000000 |
| enriched | regulation of cellular component organization | 74 | 32.883700 | 0.000000 |
| enriched | regulation of protein phosphorylation | 53 | 19.203600 | 0.000000 |
| enriched | positive regulation of nitrogen compound metabolic process | 86 | 42.397300 | 0.000000 |
| enriched | regulation of programmed cell death | 55 | 20.913600 | 0.000000 |
| enriched | positive regulation of protein serine/threonine kinase activity | 24 | 4.505730 | 0.000000 |
| enriched | cellular response to organic substance | 51 | 18.687900 | 0.000000 |
| enriched | regulation of phosphorus metabolic process | 59 | 23.790800 | 0.000000 |
| enriched | regulation of cellular process | 194 | 142.148000 | 0.000000 |
| enriched | regulation of signal transduction | 84 | 41.637300 | 0.000000 |
| enriched | positive regulation of catalytic activity | 51 | 19.054300 | 0.000000 |
| enriched | regulation of nitrogen compound metabolic process | 126 | 76.543100 | 0.000000 |
| enriched | regulation of intracellular signal transduction | 59 | 24.374400 | 0.000000 |
| enriched | regulation of metabolic process | 138 | 87.590200 | 0.000000 |
| enriched | regulation of protein metabolic process | 77 | 37.063700 | 0.000000 |
| enriched | regulation of cell death | 56 | 22.501500 | 0.000000 |
| enriched | regulation of apoptotic process | 53 | 20.642200 | 0.000000 |
| enriched | response to organonitrogen compound | 37 | 11.155700 | 0.000000 |
| enriched | regulation of MAP kinase activity | 23 | 4.532870 | 0.000000 |
| enriched | regulation of cell population proliferation | 54 | 21.510800 | 0.000000 |
| enriched | regulation of neuron differentiation | 32 | 8.767170 | 0.000000 |
| enriched | regulation of cellular protein metabolic process | 73 | 34.824400 | 0.000000 |

| | | | | |
|---|---|---|---|---|
| enriched | regulation of neurogenesis | 36 | 10.925000 | 0.000000 |
| enriched | response to nitrogen compound | 38 | 12.024300 | 0.000000 |
| enriched | positive regulation of MAP kinase activity | 20 | 3.487870 | 0.000000 |
| enriched | regulation of MAPK cascade | 34 | 9.975030 | 0.000000 |
| enriched | cellular process | 230 | 186.798000 | 0.000000 |
| enriched | regulation of nervous system development | 38 | 12.295700 | 0.000000 |
| enriched | positive regulation of intracellular signal transduction | 40 | 13.517200 | 0.000000 |
| enriched | response to peptide | 23 | 5.102870 | 0.000000 |
| enriched | biological process | 259 | 229.222000 | 0.000000 |
| enriched | positive regulation of MAPK cascade | 27 | 7.192880 | 0.000000 |
| enriched | negative regulation of cell death | 38 | 13.205000 | 0.000000 |
| enriched | positive regulation of cellular protein metabolic process | 51 | 21.347900 | 0.000000 |
| enriched | negative regulation of programmed cell death | 36 | 12.065000 | 0.000000 |
| enriched | phosphorylation | 42 | 15.620800 | 0.000000 |
| enriched | positive regulation of protein metabolic process | 53 | 22.732200 | 0.000000 |
| enriched | signal transduction | 106 | 63.880900 | 0.000000 |
| enriched | regulation of cell cycle G1/S phase transition | 15 | 2.225720 | 0.000000 |
| enriched | positive regulation of signal transduction | 52 | 22.365800 | 0.000000 |
| enriched | developmental process | 110 | 67.830200 | 0.000000 |
| enriched | negative regulation of apoptotic process | 35 | 11.861500 | 0.000000 |
| enriched | negative regulation of signaling | 45 | 17.982200 | 0.000000 |
| enriched | cellular protein modification process | 76 | 40.293700 | 0.000000 |
| enriched | negative regulation of G1/S transition of mitotic cell cycle | 12 | 1.384290 | 0.000000 |
| enriched | protein modification process | 76 | 40.293700 | 0.000000 |
| enriched | positive regulation of protein kinase activity | 26 | 7.125020 | 0.000000 |
| enriched | regulation of G1/S transition of mitotic cell cycle | 14 | 2.022150 | 0.000001 |
| enriched | positive regulation of kinase activity | 27 | 7.695020 | 0.000001 |
| enriched | regulation of cellular component movement | 37 | 13.286500 | 0.000001 |
| enriched | regulation of neuron projection development | 25 | 6.704300 | 0.000001 |
| enriched | negative regulation of cell cycle G1/S phase transition | 12 | 1.425000 | 0.000001 |
| enriched | negative regulation of signal transduction | 42 | 16.625000 | 0.000001 |
| enriched | positive regulation of phosphorylation | 38 | 14.127900 | 0.000001 |
| enriched | protein phosphorylation | 35 | 12.512900 | 0.000001 |
| enriched | cell surface receptor signaling pathway | 60 | 29.192200 | 0.000001 |
| enriched | regulation of cell migration | 33 | 11.413600 | 0.000001 |
| enriched | cellular protein metabolic process | 80 | 44.555100 | 0.000001 |
| enriched | anatomical structure morphogenesis | 45 | 19.081500 | 0.000002 |
| enriched | positive regulation of cell population proliferation | 34 | 12.214300 | 0.000002 |
| enriched | positive regulation of protein phosphorylation | 36 | 13.435700 | 0.000002 |
| enriched | regulation of cell motility | 34 | 12.282200 | 0.000002 |
| enriched | positive regulation of cell development | 25 | 7.342160 | 0.000003 |
| enriched | regulation of plasma membrane bounded cell projection organization | 28 | 8.997880 | 0.000003 |
| enriched | animal organ development | 41 | 16.977900 | 0.000003 |
| enriched | regulation of organelle organization | 41 | 16.977900 | 0.000003 |
| enriched | regulation of locomotion | 35 | 13.218600 | 0.000004 |
| enriched | regulation of cell projection organization | 28 | 9.120020 | 0.000004 |
| enriched | regulation of transcription by RNA polymerase II | 60 | 30.386500 | 0.000004 |
| enriched | response to growth factor | 18 | 4.125730 | 0.000005 |
| enriched | negative regulation of cell differentiation | 28 | 9.418600 | 0.000007 |
| enriched | positive regulation of cellular biosynthetic process | 54 | 26.572900 | 0.000008 |
| enriched | positive regulation of neurogenesis | 22 | 6.351450 | 0.000011 |
| enriched | cellular response to organonitrogen compound | 21 | 5.849300 | 0.000011 |
| enriched | peptidyl-serine phosphorylation | 13 | 2.280010 | 0.000012 |
| enriched | regulation of cellular response to stress | 28 | 9.730740 | 0.000013 |
| enriched | response to peptide hormone | 17 | 3.976440 | 0.000013 |
| enriched | positive regulation of apoptotic process | 26 | 8.617880 | 0.000013 |

| | | | | |
|---|---|---|---|---|
| enriched | positive regulation of programmed cell death | 26 | 8.685740 | 0.000015 |
| enriched | positive regulation of cell death | 27 | 9.323600 | 0.000017 |
| enriched | protein metabolic process | 87 | 53.430900 | 0.000017 |
| enriched | positive regulation of macromolecule biosynthetic process | 51 | 25.392200 | 0.000022 |
| enriched | response to stimulus | 109 | 72.838100 | 0.000022 |
| enriched | negative regulation of mitotic cell cycle phase transition | 14 | 2.850010 | 0.000024 |
| enriched | positive regulation of protein modification process | 38 | 16.407900 | 0.000024 |
| enriched | anatomical structure development | 76 | 44.921500 | 0.000026 |
| enriched | negative regulation of cellular metabolic process | 62 | 33.847200 | 0.000026 |
| enriched | positive regulation of cell migration | 22 | 6.772160 | 0.000027 |
| enriched | positive regulation of cellular component movement | 23 | 7.315020 | 0.000027 |
| enriched | vesicle-mediated transport | 46 | 22.026500 | 0.000027 |
| enriched | regulation of RNA metabolic process | 81 | 49.332300 | 0.000032 |
| enriched | positive regulation of gene expression | 52 | 26.695100 | 0.000038 |
| enriched | positive regulation of locomotion | 23 | 7.491450 | 0.000039 |
| enriched | transport | 89 | 56.389400 | 0.000041 |
| enriched | phosphate-containing compound metabolic process | 52 | 26.803600 | 0.000042 |
| enriched | thymus development | 7 | 0.610716 | 0.000042 |
| enriched | negative regulation of cell cycle | 23 | 7.545730 | 0.000042 |
| enriched | response to cytokine | 23 | 7.640730 | 0.000052 |
| enriched | response to mechanical stimulus | 13 | 2.660010 | 0.000052 |
| enriched | positive regulation of cell motility | 22 | 7.111450 | 0.000054 |
| enriched | positive regulation of neuron differentiation | 18 | 4.994300 | 0.000055 |
| enriched | regulation of nucleic acid-templated transcription | 76 | 45.980100 | 0.000055 |
| enriched | cytokine-mediated signaling pathway | 25 | 8.848590 | 0.000056 |
| enriched | regulation of RNA biosynthetic process | 76 | 46.048000 | 0.000056 |
| enriched | regulation of transcription, DNA-templated | 75 | 45.220100 | 0.000056 |
| enriched | regulation of epithelial cell differentiation | 11 | 1.927150 | 0.000065 |
| enriched | cell differentiation | 49 | 25.107200 | 0.000066 |
| enriched | cell migration | 29 | 11.386500 | 0.000066 |
| enriched | negative regulation of protein metabolic process | 34 | 14.602900 | 0.000066 |
| enriched | leukocyte migration | 16 | 4.098580 | 0.000066 |
| enriched | response to organic cyclic compound | 27 | 10.192200 | 0.000069 |
| enriched | response to hypoxia | 15 | 3.664300 | 0.000073 |
| enriched | reproductive structure development | 15 | 3.691440 | 0.000079 |
| enriched | regulation of cyclin-dependent protein serine/threonine kinase activity | 9 | 1.302860 | 0.000099 |
| enriched | positive regulation of cell differentiation | 31 | 12.960700 | 0.000100 |
| enriched | positive regulation of nucleic acid-templated transcription | 44 | 21.890800 | 0.000100 |
| enriched | positive regulation of RNA biosynthetic process | 44 | 21.904300 | 0.000100 |
| enriched | regulation of dendrite development | 11 | 2.049290 | 0.000106 |
| enriched | regulation of mitochondrion organization | 12 | 2.456440 | 0.000106 |
| enriched | response to hormone | 25 | 9.255740 | 0.000106 |
| enriched | negative regulation of cellular protein metabolic process | 32 | 13.707200 | 0.000111 |
| enriched | response to reactive oxygen species | 12 | 2.470010 | 0.000111 |
| enriched | regulation of gene expression | 89 | 58.126600 | 0.000115 |
| enriched | odontogenesis of dentin-containing tooth | 8 | 1.017860 | 0.000115 |
| enriched | aging | 14 | 3.379290 | 0.000120 |
| enriched | positive regulation of transcription, DNA-templated | 42 | 20.710100 | 0.000125 |
| enriched | chemokine (C-X-C motif) ligand 12 signaling pathway | 3 | 0.054286 | 0.000138 |
| enriched | post-embryonic camera-type eye development | 3 | 0.054286 | 0.000138 |
| enriched | positive regulation of organelle organization | 23 | 8.237880 | 0.000138 |
| enriched | neuron projection development | 14 | 3.433580 | 0.000139 |
| enriched | cell motility | 30 | 12.607900 | 0.000141 |
| enriched | regulation of epithelial cell proliferation | 16 | 4.451440 | 0.000157 |
| enriched | regulation of cell growth | 18 | 5.510010 | 0.000163 |
| enriched | negative regulation of protein serine/threonine kinase activity | 10 | 1.791430 | 0.000176 |

| | | | | |
|---|---|---|---|---|
| enriched | macromolecule metabolic process | 112 | 79.379500 | 0.000180 |
| enriched | cell activation | 31 | 13.462900 | 0.000184 |
| enriched | negative regulation of intracellular signal transduction | 20 | 6.704300 | 0.000196 |
| enriched | regulation of smooth muscle cell proliferation | 10 | 1.818580 | 0.000196 |
| enriched | odontogenesis | 9 | 1.452150 | 0.000200 |
| enriched | transmembrane receptor protein tyrosine kinase signaling pathway | 20 | 6.758590 | 0.000216 |
| enriched | negative regulation of mitotic cell cycle | 15 | 4.085010 | 0.000216 |
| enriched | negative regulation of transcription by RNA polymerase II | 28 | 11.657900 | 0.000223 |
| enriched | leukocyte apoptotic process | 5 | 0.339287 | 0.000239 |
| enriched | cellular response to growth factor stimulus | 14 | 3.650720 | 0.000251 |
| enriched | regulation of cell cycle | 34 | 15.824300 | 0.000273 |
| enriched | regulation of lymphocyte migration | 7 | 0.855002 | 0.000282 |
| enriched | animal organ morphogenesis | 20 | 6.948590 | 0.000307 |
| enriched | locomotion | 31 | 13.883600 | 0.000307 |
| enriched | response to inorganic substance | 20 | 6.962160 | 0.000311 |
| enriched | cell death | 27 | 11.291500 | 0.000325 |
| enriched | regulation of T cell migration | 6 | 0.597144 | 0.000326 |
| enriched | cellular response to reactive oxygen species | 9 | 1.560720 | 0.000329 |
| enriched | gland development | 13 | 3.284290 | 0.000338 |
| enriched | negative regulation of cell cycle process | 15 | 4.275010 | 0.000338 |
| enriched | regulation of cysteine-type endopeptidase activity involved in apoptotic process | 12 | 2.822860 | 0.000338 |
| enriched | apoptotic process | 23 | 8.835020 | 0.000343 |
| enriched | positive regulation of mitochondrion organization | 9 | 1.574290 | 0.000343 |
| enriched | regulation of establishment of endothelial barrier | 4 | 0.190001 | 0.000343 |
| enriched | reproductive process | 38 | 18.891500 | 0.000343 |
| enriched | vasculogenesis | 7 | 0.895717 | 0.000352 |
| enriched | cell projection organization | 26 | 10.762200 | 0.000362 |
| enriched | immune system process | 51 | 28.676500 | 0.000362 |
| enriched | negative regulation of gene expression | 43 | 22.650800 | 0.000373 |
| enriched | positive regulation of neuron projection development | 14 | 3.827150 | 0.000373 |
| enriched | regulation of growth | 23 | 8.902880 | 0.000373 |
| enriched | signal transduction by protein phosphorylation | 16 | 4.872160 | 0.000382 |
| enriched | negative regulation of cyclin-dependent protein serine/threonine kinase activity | 5 | 0.393572 | 0.000442 |
| enriched | morphogenesis of an epithelium | 14 | 3.908580 | 0.000460 |
| enriched | negative regulation of cell population proliferation | 23 | 9.106450 | 0.000513 |
| enriched | regulation of cell adhesion | 23 | 9.106450 | 0.000513 |
| enriched | negative regulation of cyclin-dependent protein kinase activity | 5 | 0.407144 | 0.000513 |
| enriched | response to drug | 25 | 10.477200 | 0.000595 |
| enriched | negative regulation of catalytic activity | 25 | 10.531500 | 0.000643 |
| enriched | regulation of DNA-binding transcription factor activity | 17 | 5.686440 | 0.000674 |
| enriched | cell-matrix adhesion | 8 | 1.357150 | 0.000686 |
| enriched | phosphatidylinositol biosynthetic process | 8 | 1.357150 | 0.000686 |
| enriched | tissue morphogenesis | 16 | 5.157160 | 0.000706 |
| enriched | regulation of leukocyte migration | 11 | 2.619290 | 0.000712 |
| enriched | nitrogen compound metabolic process | 120 | 89.721000 | 0.000745 |
| enriched | positive regulation of transcription by RNA polymerase II | 33 | 16.109300 | 0.000751 |
| enriched | platelet activation | 8 | 1.384290 | 0.000769 |
| enriched | phosphatidylinositol 3-kinase signaling | 5 | 0.447858 | 0.000776 |
| enriched | response to fluid shear stress | 5 | 0.447858 | 0.000776 |
| enriched | regulation of cysteine-type endopeptidase activity | 12 | 3.135010 | 0.000795 |
| enriched | exocytosis | 24 | 10.070000 | 0.000800 |
| enriched | post-embryonic animal organ development | 3 | 0.095000 | 0.000823 |
| enriched | response to toxic substance | 16 | 5.252160 | 0.000830 |
| enriched | heart development | 11 | 2.687150 | 0.000851 |

| | | | | |
|---|---|---|---|---|
| enriched | regulation of GTPase activity | 18 | 6.419300 | 0.000862 |
| enriched | programmed cell death | 25 | 10.789300 | 0.000864 |
| enriched | cellular response to oxidative stress | 11 | 2.727860 | 0.000958 |
| enriched | positive regulation of mitochondrial outer membrane permeabilization involved in apoptotic signaling pathway | 5 | 0.475001 | 0.000992 |
| enriched | cellular response to cytokine stimulus | 17 | 5.930730 | 0.001019 |
| enriched | developmental process involved in reproduction | 27 | 12.295700 | 0.001072 |
| enriched | negative regulation of protein phosphorylation | 16 | 5.387870 | 0.001072 |
| enriched | negative regulation of cysteine-type endopeptidase activity involved in apoptotic process | 7 | 1.099290 | 0.001086 |
| enriched | regulation of cell cycle phase transition | 17 | 5.971440 | 0.001086 |
| enriched | hematopoietic or lymphoid organ development | 10 | 2.320720 | 0.001111 |
| enriched | regulated exocytosis | 22 | 9.052170 | 0.001127 |
| enriched | Ras protein signal transduction | 12 | 3.284290 | 0.001141 |
| enriched | secretion by cell | 28 | 13.055700 | 0.001152 |
| enriched | positive regulation of GTPase activity | 16 | 5.442160 | 0.001163 |
| enriched | B cell apoptotic process | 3 | 0.108572 | 0.001183 |
| enriched | cellular response to insulin-like growth factor stimulus | 3 | 0.108572 | 0.001183 |
| enriched | positive regulation of cell proliferation involved in kidney development | 3 | 0.108572 | 0.001183 |
| enriched | post-embryonic animal organ morphogenesis | 3 | 0.108572 | 0.001183 |
| enriched | enzyme linked receptor protein signaling pathway | 22 | 9.120020 | 0.001208 |
| enriched | positive regulation of apoptotic signaling pathway | 10 | 2.361430 | 0.001223 |
| enriched | activation of protein kinase activity | 14 | 4.383580 | 0.001268 |
| enriched | negative regulation of cell development | 14 | 4.397150 | 0.001304 |
| enriched | positive regulation of transport | 28 | 13.191500 | 0.001307 |
| enriched | regulation of mitotic cell cycle phase transition | 16 | 5.523590 | 0.001317 |
| enriched | regulation of autophagy | 14 | 4.410730 | 0.001328 |
| enriched | regulation of cell migration involved in sprouting angiogenesis | 5 | 0.515716 | 0.001332 |
| enriched | secretion | 30 | 14.616500 | 0.001332 |
| enriched | response to antibiotic | 13 | 3.908580 | 0.001424 |
| enriched | MAPK cascade | 14 | 4.451440 | 0.001435 |
| enriched | positive regulation of smooth muscle cell proliferation | 7 | 1.167150 | 0.001439 |
| enriched | cardiocyte differentiation | 5 | 0.529287 | 0.001480 |
| enriched | response to hydrogen peroxide | 8 | 1.560720 | 0.001480 |
| enriched | helper T cell diapedesis | 2 | 0.027143 | 0.001516 |
| enriched | hepatic immune response | 2 | 0.027143 | 0.001516 |
| enriched | lymph vessel morphogenesis | 2 | 0.027143 | 0.001516 |
| enriched | negative regulation of creatine transmembrane transporter activity | 2 | 0.027143 | 0.001516 |
| enriched | response to ultrasound | 2 | 0.027143 | 0.001516 |
| enriched | regulation of epithelial cell migration | 11 | 2.931440 | 0.001537 |
| enriched | negative regulation of biosynthetic process | 38 | 20.655800 | 0.001573 |
| enriched | coronary artery morphogenesis | 3 | 0.122143 | 0.001587 |
| enriched | mannose metabolic process | 3 | 0.122143 | 0.001587 |
| enriched | cellular localization | 49 | 29.205800 | 0.001594 |
| enriched | positive regulation of protein catabolic process | 11 | 2.958580 | 0.001631 |
| enriched | negative regulation of cysteine-type endopeptidase activity | 7 | 1.207860 | 0.001673 |
| enriched | signal transduction by p53 class mediator | 8 | 1.601430 | 0.001673 |
| enriched | regulation of binding | 15 | 5.116440 | 0.001689 |
| enriched | sensory organ development | 8 | 1.628580 | 0.001862 |
| enriched | negative regulation of protein kinase activity | 11 | 3.012870 | 0.001862 |
| enriched | response to tumor necrosis factor | 9 | 2.062860 | 0.001862 |
| enriched | endoplasmic reticulum calcium ion homeostasis | 4 | 0.312144 | 0.001876 |
| enriched | positive regulation of cell projection organization | 15 | 5.184300 | 0.001906 |
| enriched | behavior | 19 | 7.600020 | 0.001929 |
| enriched | developmental growth | 13 | 4.085010 | 0.001966 |

| | | | | |
|---|---|---|---|---|
| enriched | positive regulation of hydrolase activity | 23 | 10.205700 | 0.001966 |
| enriched | cell-substrate adhesion | 9 | 2.103580 | 0.002101 |
| enriched | hypotonic response | 3 | 0.135715 | 0.002104 |
| enriched | negative regulation of cell size | 3 | 0.135715 | 0.002104 |
| enriched | endocrine pancreas development | 4 | 0.325715 | 0.002158 |
| enriched | cellular response to peptide | 11 | 3.080720 | 0.002167 |
| enriched | cellular metabolic process | 127 | 99.139500 | 0.002231 |
| enriched | negative regulation of phosphorylation | 16 | 5.890020 | 0.002321 |
| enriched | negative regulation of intrinsic apoptotic signaling pathway | 7 | 1.289290 | 0.002332 |
| enriched | regulation of immune system process | 37 | 20.411500 | 0.002332 |
| enriched | phosphatidylinositol metabolic process | 9 | 2.144290 | 0.002339 |
| enriched | intrinsic apoptotic signaling pathway in response to DNA damage | 6 | 0.922860 | 0.002351 |
| enriched | regulation of mitochondrial membrane permeability | 6 | 0.922860 | 0.002351 |
| enriched | metabolic process | 137 | 109.020000 | 0.002409 |
| enriched | intracellular transport | 36 | 19.719300 | 0.002425 |
| enriched | response to extracellular stimulus | 17 | 6.541450 | 0.002453 |
| enriched | positive regulation of cell adhesion | 15 | 5.347160 | 0.002459 |
| enriched | regulation of intrinsic apoptotic signaling pathway | 9 | 2.171430 | 0.002499 |
| enriched | regulation of mitochondrial outer membrane permeabilization involved in apoptotic signaling pathway | 5 | 0.610716 | 0.002500 |
| enriched | small GTPase mediated signal transduction | 13 | 4.220730 | 0.002500 |
| enriched | anterior/posterior pattern specification | 9 | 2.185010 | 0.002578 |
| enriched | tube morphogenesis | 12 | 3.691440 | 0.002578 |
| enriched | peptidyl-threonine phosphorylation | 6 | 0.950003 | 0.002650 |
| enriched | response to oxidative stress | 14 | 4.817870 | 0.002650 |
| enriched | kidney development | 8 | 1.750720 | 0.002691 |
| enriched | leukocyte activation | 25 | 11.875000 | 0.002691 |
| enriched | chemical homeostasis | 29 | 14.711500 | 0.002706 |
| enriched | regulation of cell morphogenesis | 17 | 6.622870 | 0.002706 |
| enriched | positive regulation of cell growth | 9 | 2.225720 | 0.002878 |
| enriched | regulation of vasculature development | 13 | 4.315730 | 0.002978 |
| enriched | regulation of protein stability | 12 | 3.772870 | 0.003030 |
| enriched | negative regulation of kinase activity | 11 | 3.270720 | 0.003267 |
| enriched | coronary vasculature morphogenesis | 3 | 0.162858 | 0.003371 |
| enriched | positive regulation of keratinocyte migration | 3 | 0.162858 | 0.003371 |
| enriched | regulation of IRE1-mediated unfolded protein response | 3 | 0.162858 | 0.003371 |
| enriched | leukocyte degranulation | 17 | 6.772160 | 0.003373 |
| enriched | positive regulation of immune system process | 27 | 13.503600 | 0.003373 |
| enriched | response to nutrient levels | 16 | 6.161440 | 0.003404 |
| enriched | regulation of cellular localization | 23 | 10.721500 | 0.003406 |
| enriched | cell chemotaxis | 10 | 2.782150 | 0.003411 |
| enriched | cell adhesion | 25 | 12.119300 | 0.003439 |
| enriched | ERK1 and ERK2 cascade | 4 | 0.380001 | 0.003469 |
| enriched | T cell migration | 4 | 0.380001 | 0.003469 |
| enriched | regulation of developmental growth | 13 | 4.410730 | 0.003482 |
| enriched | PMA-inducible membrane protein ectodomain proteolysis | 2 | 0.040714 | 0.003518 |
| enriched | cerebellar neuron development | 2 | 0.040714 | 0.003518 |
| enriched | extracellular polysaccharide biosynthetic process | 2 | 0.040714 | 0.003518 |
| enriched | fibroblast apoptotic process | 2 | 0.040714 | 0.003518 |
| enriched | neurofilament bundle assembly | 2 | 0.040714 | 0.003518 |
| enriched | regulation of dendritic spine development | 6 | 1.017860 | 0.003518 |
| enriched | protein localization | 44 | 26.505100 | 0.003529 |
| enriched | chemotaxis | 13 | 4.451440 | 0.003669 |
| enriched | positive regulation of peptidyl-serine phosphorylation | 7 | 1.425000 | 0.003669 |
| enriched | regulation of peptidyl-serine phosphorylation | 8 | 1.859290 | 0.003669 |
| enriched | regulation of protein localization | 27 | 13.639300 | 0.003694 |

| | | | | |
|---|---|---|---|---|
| enriched | striated muscle cell differentiation | 5 | 0.692145 | 0.003957 |
| enriched | morphogenesis of a branching epithelium | 8 | 1.886430 | 0.003981 |
| enriched | homeostatic process | 36 | 20.438600 | 0.004055 |
| enriched | DNA damage response, signal transduction by p53 class mediator | 6 | 1.058570 | 0.004184 |
| enriched | negative regulation of myeloid cell differentiation | 6 | 1.058570 | 0.004184 |
| enriched | negative regulation of transcription, DNA-templated | 30 | 15.960000 | 0.004186 |
| enriched | macromolecule localization | 44 | 26.776500 | 0.004209 |
| enriched | activation of MAPKK activity | 5 | 0.705716 | 0.004233 |
| enriched | positive regulation of release of cytochrome c from mitochondria | 4 | 0.407144 | 0.004243 |
| enriched | regulation of proteolysis | 21 | 9.608600 | 0.004318 |
| enriched | anatomical structure formation involved in morphogenesis | 21 | 9.622170 | 0.004378 |
| enriched | regulation of membrane permeability | 6 | 1.072150 | 0.004378 |
| enriched | immune response-regulating cell surface receptor signaling pathway | 13 | 4.573580 | 0.004509 |
| enriched | cell cycle arrest | 8 | 1.940720 | 0.004617 |
| enriched | response to lipid | 22 | 10.355000 | 0.004620 |
| enriched | extrinsic apoptotic signaling pathway in absence of ligand | 4 | 0.420715 | 0.004686 |
| enriched | face morphogenesis | 4 | 0.420715 | 0.004686 |
| enriched | negative regulation of intrinsic apoptotic signaling pathway in response to DNA damage | 4 | 0.420715 | 0.004686 |
| enriched | intrinsic apoptotic signaling pathway | 8 | 1.954290 | 0.004758 |
| enriched | Fc-epsilon receptor signaling pathway | 7 | 1.506430 | 0.004762 |
| enriched | cellular response to antibiotic | 7 | 1.506430 | 0.004762 |
| enriched | learning or memory | 11 | 3.487870 | 0.004794 |
| enriched | regulation of protein catabolic process | 14 | 5.211440 | 0.004794 |
| enriched | response to glucocorticoid | 8 | 1.967860 | 0.004903 |

#### GO_-_Molecular_Function

| Type | Name | #Hits | Expected score | p-Value |
|---|---|---|---|---|
| enriched | protein binding | 209 | 158.365000 | 0.000000 |
| enriched | kinase binding | 32 | 9.880030 | 0.000001 |
| enriched | protein kinase binding | 30 | 8.767170 | 0.000001 |
| enriched | enzyme binding | 57 | 29.477200 | 0.000058 |
| enriched | protein-containing complex binding | 33 | 14.616500 | 0.000687 |
| enriched | ATPase inhibitor activity | 3 | 0.067857 | 0.000972 |
| enriched | molecular function | 251 | 228.869000 | 0.000972 |
| enriched | molecular function regulator | 45 | 23.709300 | 0.000972 |
| enriched | protein phosphorylated amino acid binding | 6 | 0.692145 | 0.002274 |
| enriched | DNA-binding transcription factor activity, RNA polymerase II-specific | 31 | 14.779300 | 0.002347 |
| enriched | transcription regulatory region DNA binding | 28 | 12.689300 | 0.002347 |
| enriched | double-stranded DNA binding | 28 | 13.001500 | 0.002815 |
| enriched | enzyme regulator activity | 29 | 13.666500 | 0.002815 |
| enriched | sequence-specific DNA binding | 32 | 15.837900 | 0.002815 |
| enriched | proximal promoter sequence-specific DNA binding | 19 | 7.301450 | 0.002998 |
| enriched | transcription regulatory region sequence-specific DNA binding | 25 | 11.142200 | 0.002998 |
| enriched | phosphomannomutase activity | 2 | 0.027143 | 0.003465 |
| enriched | phosphotyrosine residue binding | 5 | 0.542859 | 0.003529 |
| enriched | protein serine/threonine kinase activity | 16 | 5.700020 | 0.003631 |
| enriched | phosphotransferase activity, alcohol group as acceptor | 21 | 8.943600 | 0.004428 |

#### WikiPathways

| Type | Name | #Hits | Expected score | p-Value |
|---|---|---|---|---|
| enriched | Gastrin Signaling Pathway | 18 | 1.547150 | 0.000000 |
| enriched | DNA Damage Response (only ATM dependent) | 15 | 1.492860 | 0.000000 |
| enriched | PI3K-Akt Signaling Pathway | 24 | 4.614300 | 0.000000 |
| enriched | Small cell lung cancer | 14 | 1.289290 | 0.000000 |
| enriched | Signaling Pathways in Glioblastoma | 13 | 1.112860 | 0.000000 |
| enriched | IL-5 Signaling Pathway | 9 | 0.542859 | 0.000000 |
| enriched | AGE/RAGE pathway | 10 | 0.895717 | 0.000001 |
| enriched | Hepatitis B infection | 14 | 2.049290 | 0.000001 |
| enriched | Integrated Breast Cancer Pathway | 14 | 2.062860 | 0.000001 |
| enriched | VEGFA-VEGFR2 Signaling Pathway | 17 | 3.202870 | 0.000001 |
| enriched | Epithelial to mesenchymal transition in colorectal cancer | 14 | 2.157860 | 0.000001 |
| enriched | Focal Adhesion-PI3K-Akt-mTOR-signaling pathway | 19 | 4.112150 | 0.000001 |
| enriched | T-Cell antigen Receptor (TCR) Signaling Pathway | 11 | 1.221430 | 0.000001 |
| enriched | Bladder Cancer | 8 | 0.542859 | 0.000001 |
| enriched | Chemokine signaling pathway | 14 | 2.225720 | 0.000001 |
| enriched | B Cell Receptor Signaling Pathway | 11 | 1.316430 | 0.000001 |
| enriched | PI3K-AKT-mTOR signaling pathway and therapeutic opportunities | 7 | 0.407144 | 0.000002 |
| enriched | Human Thyroid Stimulating Hormone (TSH) signaling pathway | 9 | 0.895717 | 0.000003 |
| enriched | Resistin as a regulator of inflammation | 7 | 0.447858 | 0.000003 |
| enriched | Insulin Signaling | 13 | 2.171430 | 0.000004 |
| enriched | EGF/EGFR Signaling Pathway | 13 | 2.185010 | 0.000004 |
| enriched | Regulation of Apoptosis by Parathyroid Hormone-related Protein | 6 | 0.298572 | 0.000004 |
| enriched | ErbB Signaling Pathway | 10 | 1.235000 | 0.000005 |
| enriched | IL-1 signaling pathway | 8 | 0.746431 | 0.000008 |
| enriched | Angiopoietin Like Protein 8 Regulatory Pathway | 11 | 1.791430 | 0.000019 |
| enriched | Apoptosis | 9 | 1.140000 | 0.000019 |
| enriched | Endometrial cancer | 8 | 0.855002 | 0.000019 |
| enriched | Integrated Cancer Pathway | 7 | 0.597144 | 0.000019 |
| enriched | Association Between Physico-Chemical Features and Toxicity Associated Pathways | 8 | 0.895717 | 0.000025 |
| enriched | Head and Neck Squamous Cell Carcinoma | 8 | 0.895717 | 0.000025 |
| enriched | RAC1/PAK1/p38/MMP2 Pathway | 8 | 0.922860 | 0.000030 |
| enriched | Spinal Cord Injury | 10 | 1.574290 | 0.000031 |
| enriched | Structural Pathway of Interleukin 1 (IL-1) | 7 | 0.665002 | 0.000031 |
| enriched | Photodynamic therapy-induced AP-1 survival signaling. | 7 | 0.678573 | 0.000035 |
| enriched | Photodynamic therapy-induced NF-kB survival signaling | 6 | 0.475001 | 0.000049 |
| enriched | IL-4 Signaling Pathway | 7 | 0.732859 | 0.000056 |
| enriched | Prolactin Signaling Pathway | 8 | 1.031430 | 0.000059 |
| enriched | Myometrial Relaxation and Contraction Pathways | 11 | 2.117150 | 0.000063 |
| enriched | TGF-beta Signaling Pathway | 10 | 1.791430 | 0.000084 |
| enriched | Kit receptor signaling pathway | 7 | 0.800716 | 0.000091 |
| enriched | MAPK and NFkB Signalling Pathways Inhibited by Yersinia YopJ | 4 | 0.162858 | 0.000092 |
| enriched | IL-7 Signaling Pathway | 5 | 0.339287 | 0.000109 |
| enriched | MicroRNAs in cardiomyocyte hypertrophy | 8 | 1.140000 | 0.000109 |
| enriched | IL-2 Signaling Pathway | 6 | 0.570002 | 0.000114 |
| enriched | TNF related weak inducer of apoptosis (TWEAK) Signaling Pathway | 6 | 0.570002 | 0.000114 |
| enriched | Osteopontin Signaling | 4 | 0.176429 | 0.000117 |
| enriched | IL-6 signaling pathway | 6 | 0.583573 | 0.000125 |
| enriched | Brain-Derived Neurotrophic Factor (BDNF) signaling pathway | 10 | 1.954290 | 0.000141 |
| enriched | Oncostatin M Signaling Pathway | 7 | 0.882145 | 0.000141 |
| enriched | Pancreatic adenocarcinoma pathway | 8 | 1.207860 | 0.000141 |
| enriched | Aryl Hydrocarbon Receptor Netpath | 6 | 0.624287 | 0.000168 |

| | | | | |
|---|---|---|---|---|
| enriched | Regulation of Microtubule Cytoskeleton | 6 | 0.624287 | 0.000168 |
| enriched | TNF alpha Signaling Pathway | 8 | 1.248570 | 0.000169 |
| enriched | Mammary gland development pathway - Embryonic development (Stage 1 of 4) | 4 | 0.203572 | 0.000186 |
| enriched | Fragile X Syndrome | 9 | 1.642150 | 0.000186 |
| enriched | let-7 inhibition of ES cell reprogramming | 3 | 0.081429 | 0.000213 |
| enriched | Breast cancer pathway | 10 | 2.090010 | 0.000217 |
| enriched | IL-3 Signaling Pathway | 6 | 0.665002 | 0.000217 |
| enriched | Non-small cell lung cancer | 7 | 0.977145 | 0.000230 |
| enriched | Synaptic signaling pathways associated with autism spectrum disorder | 6 | 0.678573 | 0.000236 |
| enriched | Chromosomal and microsatellite instability in colorectal cancer | 7 | 0.990717 | 0.000243 |
| enriched | Amplification and Expansion of Oncogenic Pathways as Metastatic Traits | 4 | 0.230715 | 0.000276 |
| enriched | Integrin-mediated Cell Adhesion | 8 | 1.370720 | 0.000277 |
| enriched | Wnt Signaling Pathway (Netpath) | 6 | 0.705716 | 0.000277 |
| enriched | Alpha 6 Beta 4 signaling pathway | 5 | 0.447858 | 0.000283 |
| enriched | BDNF-TrkB Signaling | 5 | 0.447858 | 0.000283 |
| enriched | Leptin signaling pathway | 7 | 1.031430 | 0.000283 |
| enriched | miRNA regulation of prostate cancer signaling pathways | 5 | 0.447858 | 0.000283 |
| enriched | Toll-like Receptor Signaling Pathway | 8 | 1.397860 | 0.000292 |
| enriched | Focal Adhesion | 11 | 2.687150 | 0.000306 |
| enriched | MFAP5 effect on permeability and motility of endothelial cells via cytoskeleton rearrangement | 4 | 0.244286 | 0.000306 |
| enriched | p38 MAPK Signaling Pathway | 5 | 0.461430 | 0.000309 |
| enriched | RANKL/RANK (Receptor activator of NFKB (ligand)) Signaling Pathway | 6 | 0.746431 | 0.000335 |
| enriched | Cell Differentiation - Index expanded | 4 | 0.257858 | 0.000368 |
| enriched | Regulation of toll-like receptor signaling pathway | 9 | 1.886430 | 0.000400 |
| enriched | Hematopoietic Stem Cell Gene Regulation by GABP alpha/beta Complex | 4 | 0.271429 | 0.000433 |
| enriched | Neovascularisation processes | 5 | 0.502144 | 0.000433 |
| enriched | Serotonin Receptor 2 and ELK-SRF/GATA4 signaling | 4 | 0.271429 | 0.000433 |
| enriched | MET in type 1 papillary renal cell carcinoma | 6 | 0.800716 | 0.000459 |
| enriched | Oxidative Damage | 5 | 0.542859 | 0.000600 |
| enriched | Role of Altered Glycolysation of MUC1 in Tumour Microenvironment | 3 | 0.122143 | 0.000600 |
| enriched | Type II diabetes mellitus | 4 | 0.298572 | 0.000600 |
| enriched | Vitamin D in inflammatory diseases | 4 | 0.298572 | 0.000600 |
| enriched | G1 to S cell cycle control | 6 | 0.868574 | 0.000679 |
| enriched | Cell Cycle | 8 | 1.628580 | 0.000686 |
| enriched | Nonalcoholic fatty liver disease | 9 | 2.062860 | 0.000686 |
| enriched | Estrogen signaling pathway | 4 | 0.312144 | 0.000686 |
| enriched | Mammary gland development pathway - Involution (Stage 4 of 4) | 3 | 0.135715 | 0.000782 |
| enriched | Angiogenesis | 4 | 0.325715 | 0.000796 |
| enriched | Interleukin-11 Signaling Pathway | 5 | 0.597144 | 0.000865 |
| enriched | Relationship between inflammation, COX-2 and EGFR | 4 | 0.339287 | 0.000917 |
| enriched | Ebola Virus Pathway on Host | 8 | 1.750720 | 0.001033 |
| enriched | Translation inhibitors in chronically activated PDGFRA cells | 5 | 0.624287 | 0.001033 |
| enriched | EPO Receptor Signaling | 4 | 0.352858 | 0.001038 |
| enriched | Thymic Stromal LymphoPoietin (TSLP) Signaling Pathway | 5 | 0.637859 | 0.001119 |
| enriched | Hepatitis C and Hepatocellular Carcinoma | 5 | 0.665002 | 0.001347 |
| enriched | MAPK Signaling Pathway | 11 | 3.338580 | 0.001451 |
| enriched | Senescence and Autophagy in Cancer | 7 | 1.425000 | 0.001463 |
| enriched | Copper homeostasis | 5 | 0.705716 | 0.001722 |
| enriched | Cardiac Progenitor Differentiation | 5 | 0.719288 | 0.001862 |

| | | | | |
|---|---|---|---|---|
| enriched | Factors and pathways affecting insulin-like growth factor (IGF1)-Akt signaling | 4 | 0.420715 | 0.001928 |
| enriched | Interferon type I signaling pathways | 5 | 0.732859 | 0.001990 |
| enriched | Cardiac Hypertrophic Response | 5 | 0.746431 | 0.002144 |
| enriched | MAPK Cascade | 4 | 0.447858 | 0.002319 |
| enriched | Signal transduction through IL1R | 4 | 0.447858 | 0.002319 |
| enriched | Viral Acute Myocarditis | 6 | 1.140000 | 0.002319 |
| enriched | miRNAs involved in DNA damage response | 3 | 0.203572 | 0.002319 |
| enriched | Oxytocin signaling | 2 | 0.054286 | 0.002493 |
| enriched | Signaling of Hepatocyte Growth Factor Receptor | 4 | 0.461430 | 0.002551 |
| enriched | Interactome of polycomb repressive complex 2 (PRC2) | 3 | 0.217143 | 0.002749 |
| enriched | Leptin Insulin Overlap | 3 | 0.230715 | 0.003275 |
| enriched | Amyotrophic lateral sclerosis (ALS) | 4 | 0.502144 | 0.003394 |
| enriched | miRNAs involvement in the immune response in sepsis | 4 | 0.502144 | 0.003394 |
| enriched | Corticotropin-releasing hormone signaling pathway | 6 | 1.248570 | 0.003473 |
| enriched | 4-hydroxytamoxifen, Dexamethasone, and Retinoic Acids Regulation of p27 Expression | 3 | 0.244286 | 0.003691 |
| enriched | Inhibition of exosome biogenesis and secretion by Manumycin A in CRPC cells | 3 | 0.244286 | 0.003691 |
| enriched | Transcription co-factors SKI and SKIL protein partners | 3 | 0.244286 | 0.003691 |
| enriched | G13 Signaling Pathway | 4 | 0.529287 | 0.003915 |
| enriched | PDGF Pathway | 4 | 0.529287 | 0.003915 |
| enriched | Pathways Affected in Adenoid Cystic Carcinoma | 5 | 0.882145 | 0.003915 |
| enriched | Extracellular vesicles in the crosstalk of cardiac cells | 3 | 0.257858 | 0.004196 |
| enriched | Neural Crest Cell Migration during Development | 4 | 0.542859 | 0.004214 |
| enriched | AMP-activated Protein Kinase (AMPK) Signaling | 5 | 0.922860 | 0.004664 |
| enriched | Imatinib and Chronic Myeloid Leukemia | 3 | 0.271429 | 0.004769 |
